# Microplastics inhibit macrophage bioenergetics impairing homeostatic function and immune responsiveness

**DOI:** 10.64898/2026.05.14.725255

**Authors:** Rajeev Dhupar, Hannah M. Udoh, Naila Noureen, Charles E. Bardawil, Xinyue Zhao, Mohsin Cheema, Shivani Tuli, Dylan Shields, Ksenia Mats, Omar Al-Bataineh, Laila Golla, Angel Wang, Ricardo H. Pineda, Melanie Koenigshoff, Shikhar Uttam, David M. Gau, Adam C. Soloff

## Abstract

Since the 1950s, micro- and nanoplastics (MNPs) have become omnipresent, representing a novel environmental hazard which continually deposits in our airways. Pulmonary macrophages (pMacs) orchestrate the balance between inflammation and tolerance required for homeostasis of the lung and are among the first immune cells to encounter inhaled MNPs. Yet, how pMacs react to plastic deposition in the lung and implications for disease remain unknown. Here, we exposed mice *in vivo*, human precision-cut lung slices (hPCLS) *ex vivo*, and monocyte-derived macrophages and cell lines to polystyrene MNPs *in vitro*. MNP deposition in the lung and extrapulmonary tissues was determined over a 1-week period and pMacs from MNP-laden lungs isolated for RNA-sequencing. We compared the effects of MNPs or diesel exhaust particulate exposures on hPCLS viability and metabolism, monocyte-derived macrophage transcription, and macrophage mitochondrial function, inflammation, and antigen presentation. MNPs readily translocated the lung and were observed in all organs examined within 1-day. pMacs from MNP-exposed mice expressed transcriptional pathways associated with endocrine system disorders, tissue remodeling, and malignant disease. Macrophage phagocytosis was impaired through decreased mitochondrial function which could be rescued pharmacologically. MNPs inhibited the ability of macrophages to effectively present OVA-antigen preventing TCR-specific activation, an effect that could be restored by blocking PD-1/PD-L1. These findings indicate that MNPs impair macrophages via unique mechanisms linking phagocytic and bioenergetic dysfunction. Loss of antigen-presenting capabilities in MNP-laden macrophages may compromise immunosurveillance. As such, MNPs have the potential to increase susceptibility to lung disease independent of the conventional mechanisms of inflammation and oxidative stress.

**Clinical relevance:**

- Bioaccumulation of micro- and nanoplastics in macrophages impairs their ability to function as antigen-presenting cells increasing susceptibility to pathogenic and malignant disease.
- Pulmonary macrophages residing in micro- and nanoplastic laden lungs possess transcriptional profiles associated with endocrine system disorders, gastrointestinal disease, and cancers.

## Introduction

Since the 1950’s, an estimated 8.3 billion tons of plastic has been generated, of which, ∼79% has accumulated in the environment^1,2^. Microplastics and nanoplastics (MNPs), plastic particles under 5mm and 1μm, respectively, have been observed in almost all tissues of the body including the respiratory tract, gastrointestinal tract, brain, and placenta^3–6^. Unlike some bacteria and fungi, human cells lack chemical processes to degrade plastics, and as such, MNPs likely bioaccumulate within tissues. Recent studies found that MNP concentrations in decedent liver and brain samples increased between 2016 and 2024 in conjunction with greater global plastic production^7^. Similarly, upon ingestion of MNPs by mice, particles accumulate in the liver, kidney, and gut over time, which was associated with decreased lipid metabolism and liver acetylcholinesterase^8^. Recent correlative studies have associated the amount of MNPs in brain tissue, the gastrointestinal tract, cervical tissue, liver, and atherosclerotic plaques with the presence or severity of diseases such as dementia, inflammatory bowel disease, and cervical cancer^5,7,9–12^. Identifying mechanisms of MNP pathobiology will be critical to inform public health and clinical interventions.

Exposure to airborne particulate matter (PM) is associated with the development and poor clinical outcomes of chronic lung disease such as bronchitis, asthma, and lung cancer. While inhaled particles greater than ∼10µm are often trapped in the upper airways and cleared via mucociliary action, small particulate matter of ≤2.5µm (PM2.5) may deposit in the alveoli of the lower airways and undergo pulmonary translocation, crossing the air-blood barrier, entering systemic circulation, and seeding distant organs^13^. Each year in the U.S., an estimated 14,000 early deaths can be attributed per each 1µg/m^3^ increase in PM2.5 exposure and an increase of 10μg/m^3^ PM2.5 raises the risk of early death by 1%^14,15^. Although the contribution of MNPs to ambient PM is subject to spatial/temporal variation and detection methods, MNPs such as degradation of textiles and tire wear particles undoubtedly account for part of its complex composition. With a lower limit of detection of ∼10 µm, studies have reported ambient indoor MNPs levels ranging from 9.5 ± 5.8 MNP/m^3^ to 1583 ± 1181 MNP/m^3^ ^16,17^. This approximates inhalation of between 105 and 17,413 particles per day at an intake of 11m^3^/day. If the particle size averaged 100µm, this would amount to inhaling approximately 0.054mm^3^ to 9.12mm^3^ of MNPs, or the equivalent of ∼600 red blood cells to a small Drosophila fruit fly each day.

Macrophages are the archetypic phagocytes of the immune system and serve at the interface between the mucosa and the environment. Pulmonary macrophages (pMacs), including alveolar, interstitial, and recruited macrophages, orchestrate the balance between inflammation and tolerance required for homeostasis of the lung. This balance is maintained through the immunologically silent clearance of apoptotic cells termed efferocytosis, the uptake, immune signaling, and presentations of pathogens via inflammatory phagocytosis, and the intracellular degradation and recycling process of autophagy^18^. Most, but not all studies indicate that macrophages respond to MNPs by generating reactive oxygen species (ROS) and inflammation^19–21^. It is possible that the inability to degrade MNPs intracellularly and subsequent “frustrated phagocytosis” underlies the onset of inflammation and potential pathobiology^22–24^. Recently, Codo et. al. demonstrated that intratracheal administration of MNPs to mice decreased efferocytosis by alveolar macrophages *in vivo* through methylglyoxal glycation of glucose-6-phosphate dehydrogenase^25^. Similarly, Merkley et. al. showed that MNPs co-localized within the autophagosome but fail to be cleared, inducing an immuno-metabolically active state^26^.

In the current study, we sought to determine how the inability to clear polystyrene MNPs impairs macrophage immune reactivity. Using human monocyte-derived macrophages (MDMs) and murine cell lines, we show that MNP-induced cytotoxicity and inflammation is highly dose-dependent. Consistent with previous reports, we found that MNPs inhibit mitochondrial function and reduce mitochondrial mass, the latter of which could be reverted with the AMP kinase activator AICAR to restore phagocytosis. Examining the crosstalk between innate and adaptive immunity, we show that MNPs suppress antigen-specific T cell receptor (TCR) activation by macrophages, in part, through upregulation of inhibitory checkpoint receptors, and that antigen-specific stimulation could be revived by blocking PD-1 and PD-L1. Using human precision-cut lung slices (hPCLS) *ex vivo* and MDMs *in vitro*, we found that MNP exposure results in greater cell damage and unique transcriptional profiles compared to treatment with diesel exhaust particulate (DEP). Lastly, following intranasal administration to mice, MNPs induced the expression of transcriptional profiles in pMacs associated with endocrine system disorders, gastrointestinal disease, and cancers. Collectively, our findings provide further insight into the mechanisms by which MNPs induced immune dysfunction.

## Materials and Methods

Detailed methods are provided in the article supplement.

### MNPs and uptake

Fluorescent (ThermoFisher) or non-fluorescent (Spherotech) polystyrene microspheres were used. MDMs were generated by culture of bead-isolated CD14^+^ cells with 20ng/ml M-CSF for 5-6 days. MDMs or Raw264.7 cells were cultured overnight in chamber slides followed by MNPs for 24hrs. Cells were fixed, permeabilized, and stained with antibodies for Rab5a, Rab7, LAMP-1, or Tom20 in permeabilization buffer. Nuclei were labeled with DAPI and microfilaments with Phalloidin. Phagocytosis of FITC-Zymosan was determined via flow cytometry with or without 1hr pre-treatment with AICAR (10-100µM).

### Cytotoxicity

Raw264.7 cells were co-cultured with MNPs for 24hrs and LDH and metabolic activity were quantified using CyQUANT-LDH and CyQUANT-XTT kits, respectively. hPCLS viability was imaged via LiveDead and ATP measured in three 4mm punches from a single hPCLS using CellTiter-Glo.

### In vitro functions

Raw264.7 cells were co-cultured with MNPs for 24hrs. Raw-Dual reporter cells were treated with DEP (25-400µg/mL), 1µm (100:1-1600:1/cell) or 4µm MNPs (10:1-160:1/cell) with or without 100ng/ml lipopolysaccharide. NF-κB activation was measured via SEAP using QUANTI-Blue and IRF activation by Lucia luciferase using QUANTI-Luc. To measure ROS, Raw264.7 cells were labeled with 500nM CellROX green and 5µm Sytox red. Cytokines were detected in 50µl culture supernatant using the Mouse Inflammation CBA. Mitochondrial membrane polarization was measured by staining with 2µM JC-1.

### Seahorse

MDMs were seeded in Seahorse microplates overnight with or without 100:1 4µm MNPs. Seahorse MitoStress Tests were performed under basal conditions and following sequential injections of oligomycin (1µM), FCCP (1µM), and rotenone/antimycin A (0.5µM each). Prior to analysis, medium was replaced with SeahorseXF Assay Medium and incubated in a non-CO₂ incubator for 1hr. Data were normalized to total cell count per well and analyzed using Wave software (Agilent).

### In vivo exposures

Female and male FVB/N mice received 3x10^7^ 1µm MNPs in sterile saline intranasally. Organs were harvested and processed without enzymatic digestion. Single-cell suspensions were immediately analyzed via flow cytometry and fluorescent MNP counts normalized to 10^6^ total events. Additional cohorts received 3x10^7^ 1µm or 5x10^5^ 4µm non-fluorescent MNPs intranasally. After 1wk lungs were harvested, digested in LiberaseTM, and pMacs isolated via flow sorting with a MA900 cell sorter.

### TCR stimulation

5x10^5^ Raw264.7 cells were cultured in 2ml DMEM overnight with 100U/ml IFNγ with or without MNPs. Unbound MNPs were removed, cells washed, then cultured with ovalbumin (OVA) protein or OVA323–339 peptide for 4hrs. DO11.10 T cells were labeled with 0.5µM CFSE and co-cultured with OVA-treated Raw264.7 cells at a 1:1 ratio for 72hrs. Non-adherent DO11.10 cells were collected, stained for viability and analyzed via flow cytometry.

### Statistical analysis

Data between groups were analyzed using non-parametric Kruskal-Wallis test followed by Bonferroni-adjusted p-values for post-hoc comparisons using and GraphPad Prism11. All *in vitro* and *ex vivo* assays represent a minimum of 3 independent experiments. For all hypothesis, statistical significance was denoted as p ≤ 0.05 (*), p ≤ 0.01 (**), and p ≤ 0.001 (***), p ≤ 0.0001 (****) with data reported as the mean ± SEM.

## Results

### The toxicity of MNPs is dose-dependent

Macrophages are known to ingest MNPs but are incapable of degrading their polymers. To assess how particle numbers affect uptake, we co-cultured Raw264.7 murine macrophages with polystyrene microspheres at increasing MNP to cell ratios. Consistent with previous reports, Raw264.7 cells readily phagocytose fluorescent MNPs of 1µm and 4µm with limited uptake of 9µm particles determined by immunofluorescence microscopy (Fig.1A). To examine retention, we treated terminally differentiated human monocyte-derived macrophages (MDMs) with 1µm MNPs and measured the number of intracellular MNPs over time. After 6 days, the number of MNPs per macrophage decreased by only 3.12%, averaging 46.5/cell compared to 48/cell detected on 1-day (Fig.1B). We next tracked phagolysosomal processing of MNPs after uptake by MDMs. Here, MDMs were co-cultured with red fluorescent 1µm MNPs and immunostained for the early phagosome (Rab5a), late phagosome (Rab7), or lysosome (LAMP-1). MNPs were predominantly retained in the early phagosome with ∼60% co-localized with Rab5 vesicles compared to ∼17.7% in late phagosomes and ∼21.8% in lysosomes (Fig.1C). As macrophages digest phagocytosed particulate via lysosomal degradation, we cultured FITC-labeled MNPs at pH 4.5. After 4hrs, there was no loss of fluorescence intensity (p = 0.841), indicating MNPs had not degraded and exposed acid-sensitive FITC molecules.

**Figure 1:**
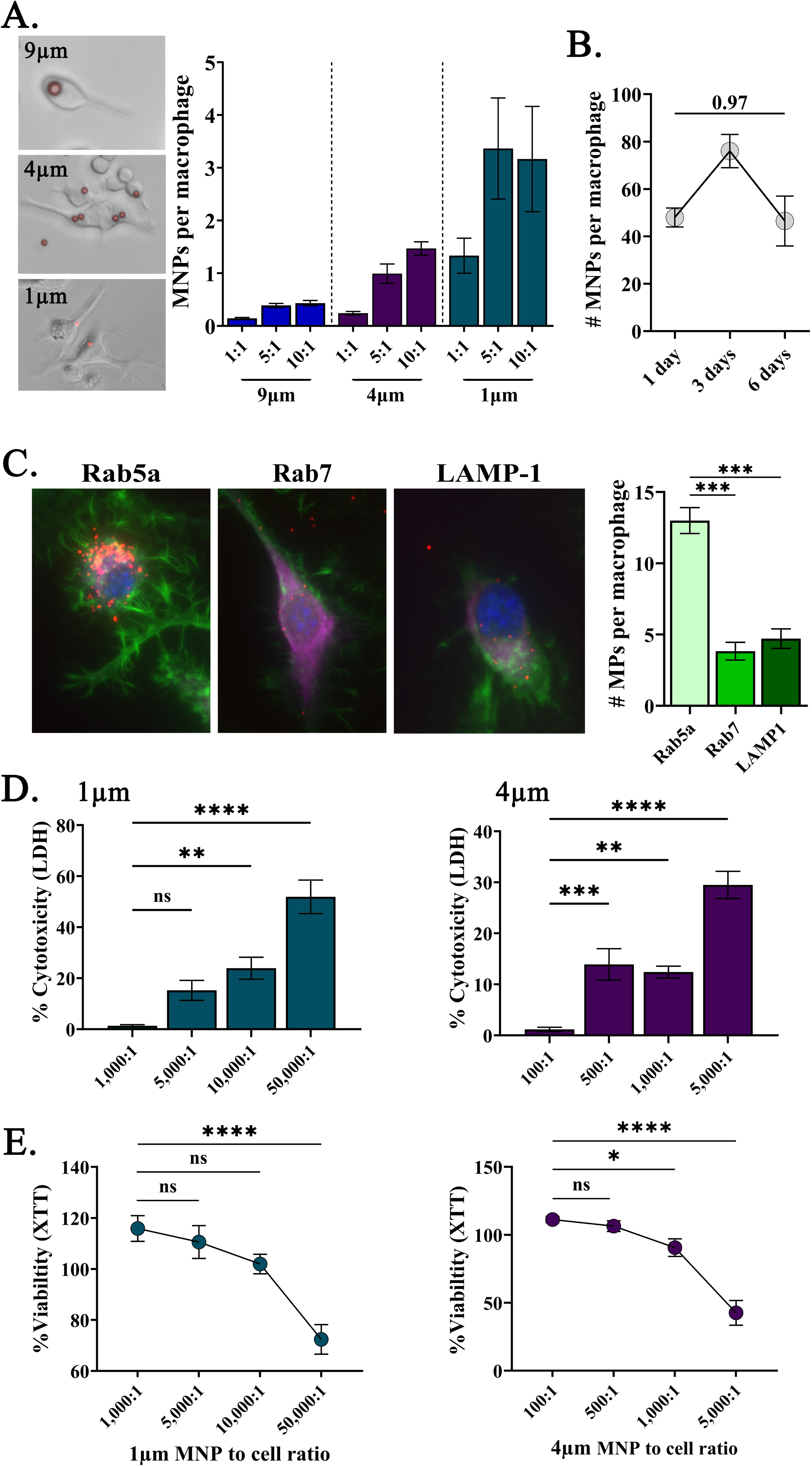
MNPs are retained in cells but only toxic at higher doses. A) Raw264.7 cells were co-cultured with 1μm, 4μm, or 9μm MNPs and uptake quantitated by fluorescence microscopy. B) Quantity of 1μm MNPs in MDMs over 6-days after 100:1 MNP/cell exposure. C) MDMs were treated with 1μm MNP (red) and stained with Rab5, Rab7, and LAMP-1 (purple), phalloidin (green), and DAPI (blue) with counts of MNPs per compartment (right). D) LDH release and E) XTT viability assays of Raw264.7 cells treated with 1μm (left) or 4μm (right) MNPs at increasing MNP to cell ratios.

MNPs have been reported to be cytotoxic, yet doses tested are frequently beyond what is likely to be experienced *in vivo*. Here, we co-cultured Raw264.7 cells for 24hrs with MNPs at ratios from 1:1 to 50,000:1 particles/cell. Significant release of LDH, indicating cell stress and/or death, was not detected until cells were treated with 10,000 1µm or 500 4µm MNPs/cell resulting in 15.25% and 13.92% LDH production compared to non-treated controls (Fig.1D).

These exposures amount to 5.5ng and 17.6ng of 1µm or 4µm MNPs per macrophage, respectively. Decreased metabolic activity by XTT assay was not observed until cells were treated with 50,000 1µm or 1,000 4µm MNPs/cell, respectively (Fig.1E). Notably, similar to Yoshioka et. al., our observed effects were associated with particle volume per cell and not concentration (Fig.S1, Table S1)^27^.

### MNPs induce differential regulation of transcriptional programs

The inability of macrophages to degrade MNPs results in frustrated phagocytosis marked by destabilization of lysosomes with protease leakage into the cytosol, autophagy, cell stress and release of damage-associated molecular patterns (DAMPs). To examine how MNPs regulate inflammation, we measured activation of the transcription factors NF-κβ and interferon response factor (IRF) family simultaneously in engineered Raw264.7 dual-reporter cells. NF-κβ is upregulated in response to inflammation, stress, and tissue injury, whereas IRFs activate following IFNα/IFNβ and JAK-STAT signaling, inducing antiviral responses and inhibiting translation. Macrophages were treated with MNPs with or without LPS to determine transcription factor activation or suppression and compared to cells administered diesel exhaust particles (DEP) as a well-characterized reference exposure^28^. In the absence of LPS, only co-culture with 1µm MNPs activated NF-κβ, whereas both 1µm and 4µm MNPs and DEP were able to increase NF-κβ above levels induced by LPS alone (Fig.2A). By contrast, treatment with either DEP or 4µm MNPs inhibited IRF activity in both the presence and absence of LPS stimulation, suggesting that they MNPs inhibit antiviral immunity while promoting inflammation (Fig.2B).

**Figure 2:**
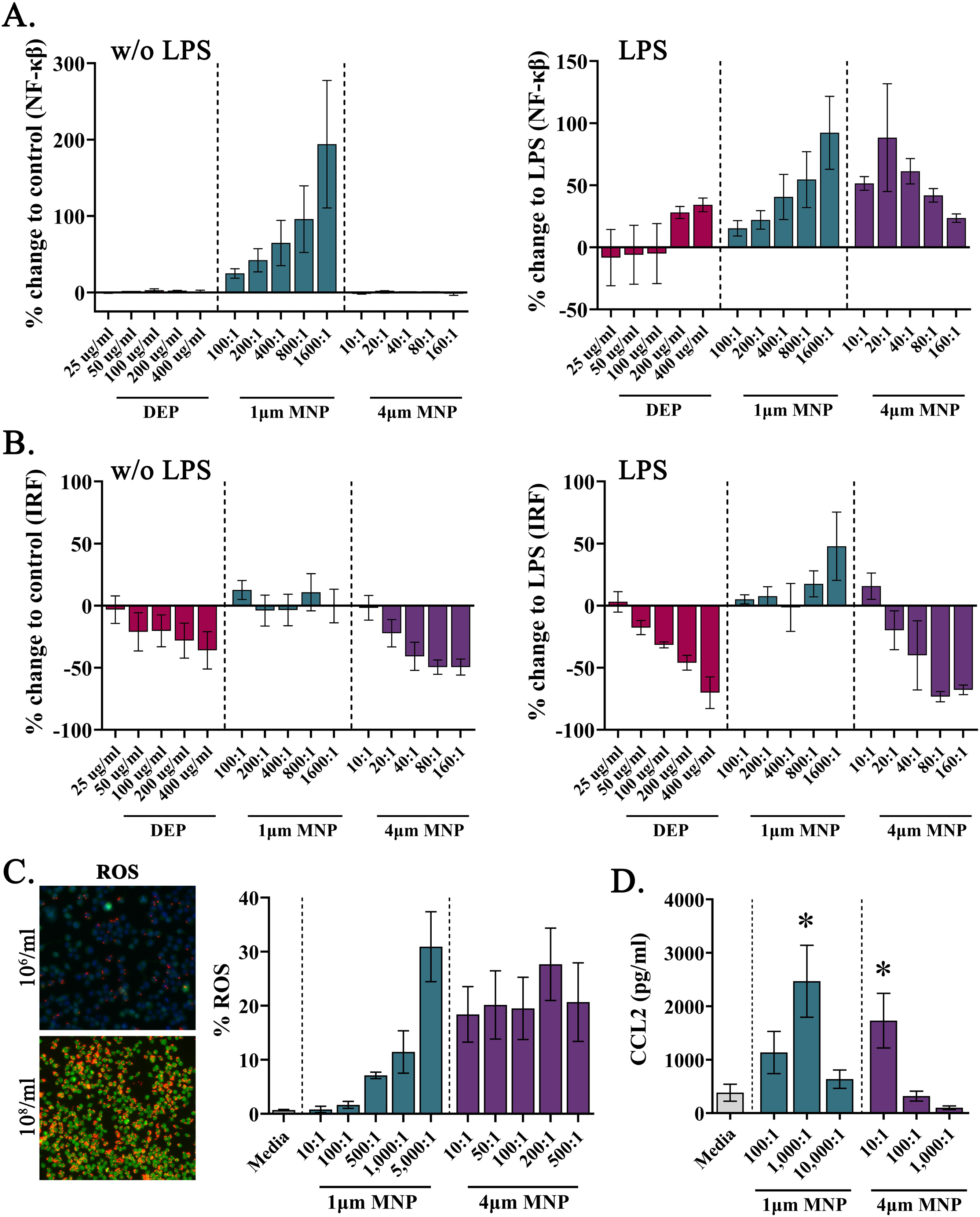
MNPs induce inflammation associated with NF-κβ activation. Dual-reporter Raw264.7 cells measuring A) NF-κβ or B) IRF activation following exposure to 1μm or 4μm MNPs in the presence (right) or absence (left) of LPS. C) The percentage of ROS positive cells by CellROX staining of Raw246.7 cells after co-culture with 1μm or 4μm MNPs. D) Levels of CCL2 in cell culture supernatant following treatment of Raw246.7 cells with 1μm or 4μm MNPs.

As reactive oxygen species (ROS) are produced following NF-κβ activation, we next measured ROS production post-exposure via CellRox assay kit. As previously reported, MNPs induced significant ROS production from Raw264.7 cells (Fig.2C) with 1µm MNPs driving a dose-dependent response whereas 4µm MNPs inducing ROS production even at lower concentrations. Notably, the levels of MNPs (particulate volume per cell) capable of inducing substantial ROS production also promote LDH release, suggesting terminal cell stress. Next, we evaluated the effect of MNPs on inflammatory cytokine production. Raw264.7 cells were co-cultured with MNPs for 24hrs and IFNγ, TNFα, IL-6, IL-12, IL-10 and CCL2 were measured by cytometric bead array. Raw264.7 cells exposed to MNPs produced significant amounts of CCL2 and, to a lesser extent, TNFα (Fig.3C and data not shown). Notably, cytokine production decreased with increasing concentrations of MNPs, again likely as a result of increased MNP-induced dysfunction and cell death.

**Figure 3:**
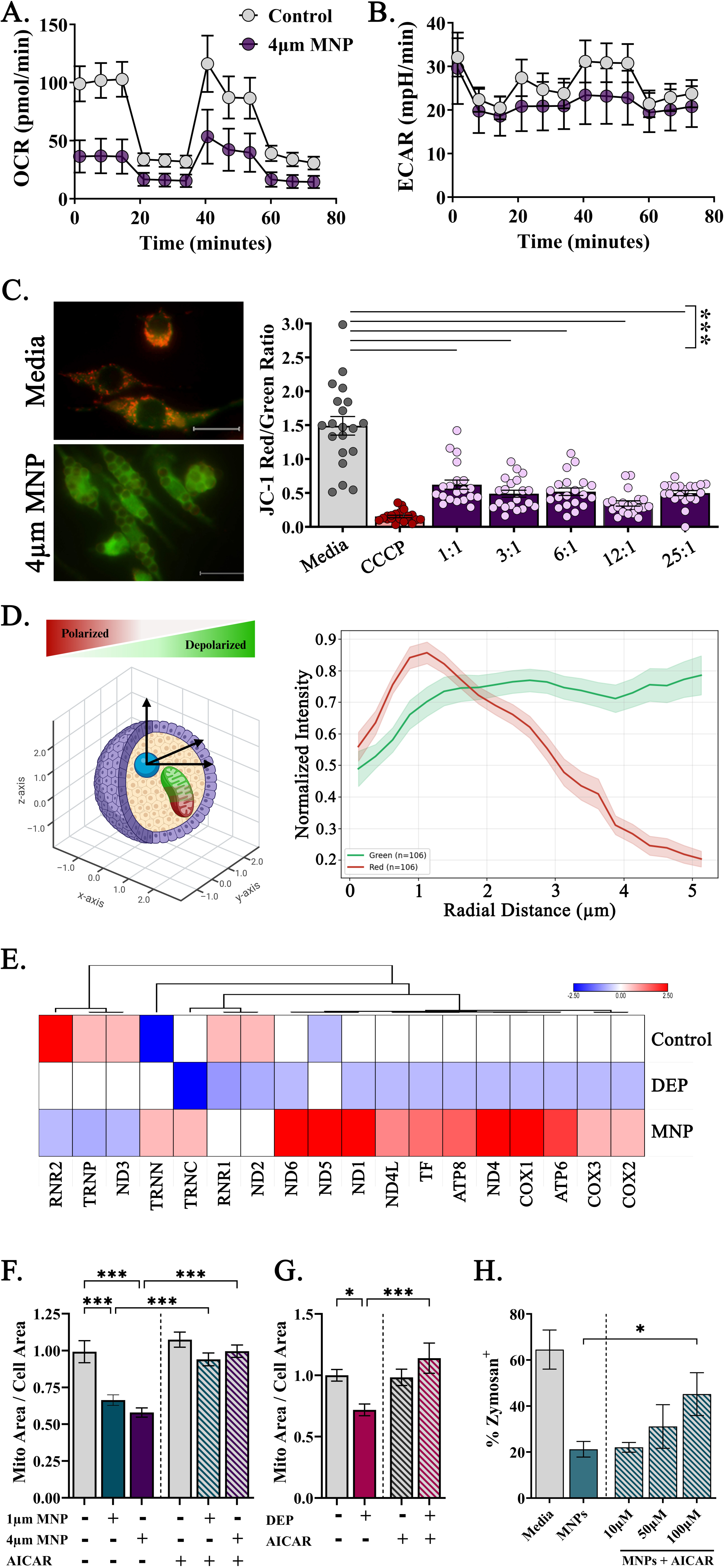
MNPs disrupt mitochondrial bioenergetics. A) Oxygen consumption rate and B) extracellular acidification rate of MDMs treated overnight with 100:1 4μm MNPs (purple) or media (grey) measured using the Seahorse XF Mito Stress Test. C) Representative images and quantitation of mitochondrial membrane polarization by JC-1 in MDMs reported as the ratio of polarized (red) over depolarized (green) per cell. D) An illustration of 3D modeling (left) and JC-1 fluorescence intensities by distance from MNP centroids (right). Values were determined from (n = 106) internalized MNP particles from confocal 3D fluorescence z-stacks of MDMs. E) Expression of mitochondrial genes from RNAseq of MDMs treated with 100:1 1μm MNPs (n = 6), 50µg/ml DEP (n = 6), or saline (n = 3) for up to 3-days. MDMs were treated with F) 100:1 1μm or 4μm MNPs or G) 50µg/ml DEP in the presence or absence of 100uM AICAR. H) MDM uptake of FITC-labeled zymosan following a 1hr pre-treatment with AICAR.

### MNPs disrupt bioenergetics and mitochondrial networks in macrophages

MNPs have been reported to bias macrophages from oxidative phosphorylation to glycolysis and decrease mitochondrial membrane polarization^26^. Here, we treated MDMs with non-fluorescent MNPs then labeled them for the Translocase of Outer Mitochondrial Membrane 20 (Tom20) receptor on the mitochondrial surface. Surprisingly, we observed the cell’s mitochondria adhering to the surface of the plastic microsphere (Fig.S2). To determine how these interactions affect macrophage respiration, we performed extracellular flux analysis using the Seahorse XF Mito Stress Test on MDMs treated at 100:1 with 4µm MNPs, a dose that induces cell stress but not death. MNP exposure significantly decreased oxygen consumption rates (OCR) indicating inhibition of oxidative phosphorylation compared to controls (Fig.3A). By contrast, MNP treatment did not significantly affect glycolysis determined via extracellular acidification rate (ECAR)(Fig.3B). Decreased OCR without a compensatory rise in ECAR suggests mitochondrial dysfunction associated with exhausted and less functional macrophages. We next used JC-1, a mitochondrial membrane potential (ΔΨm) indicator, following exposure of MDMs to 4µm MNPs. JC-1 aggregates in the mitochondrial membranes emit red fluorescence indicating high ΔΨm whereas depolarized membranes accumulate JC-1 monomers and fluoresce green indicating membrane depolarization. Administration of carbonyl cyanide m-chlorophenyl hydrazone (CCCP), a protonophore mitochondrial uncoupler, results in near complete membrane depolarization indicated by the red/green fluorescence ratio of JC-1 (0.15 ± 0.02) (Fig.3C). Notably, MNP exposures at any concentration reduced ΔΨm in MDMs compared to those cultured in media alone (Fig.3C). Using 3D spatial modelling, we measured the proximity of JC-1 red or green fluorescence with the centroid of each MNP inside macrophages (Fig.3D). Interestingly, mitochondria in close proximity (∼1µm) to a MNP particle retained high ΔΨm which was lost with increased distance from the center of the MNP (Fig.3D). We also observed a corresponding increase in green JC-1 monomers in mitochondria moving away from MNP centers, suggesting that MNPs may disrupt the organization of interconnected networks of mitochondria.

Next, we measured mitochondrial gene expression from MDMs treated with 4µm MNPs or 50µg/ml DEP *in vitro* for up to 3 days using RNAseq. Compared to saline treated controls, both MNPs and DEP reduced expression of RNR2, TRNP, ND3, RNR1, and ND2 in MDMs indicating inhibition of the mitochondrial ribosome and Complex 1 of the electron transport chain, affecting protein synthesis and ATP production, respectively (Fig.3E). Interestingly, 10 mitochondrial genes were differentially expressed in MNP vs. DEP treated MDMs relative to controls, with ND6, ND1, ND4L, TF, ATP8, ND4, COX1, ATP6, COX3, and COX2 upregulated after MNP treatment but downregulated in DEP treated MDMs (Fig.3E), suggesting potential compensation for MNP-induced damage.

We then labeled MDMs with MitoTracker prior to MNP co-culture and quantified mitochondrial mass by dividing overall mitochondrial content with cell area by confocal microscopy (Fig.3F). Compared to controls, both 1µm and 4µm MNPs (100:1) significantly decreased mitochondrial abundance, indicating impaired mitochondrial homeostasis (Fig.3F). Similarly, 50µg/ml DEP also reduced mitochondrial mass. To test if mitochondrial loss could be prevented, MDMs were administered 10-100µM 5-aminoimidazole-4-carboxamide ribonucleotide (AICAR), an AMP-activated protein kinase (AMPK) agonist known to promote mitochondrial biogenesis, 1hr before MNP/DEP administration. AICAR restored MDM mitochondrial mass to levels comparable to non-exposed controls, effectively reversing MNP/DEP-induced mitochondrial loss (Fig.3F,G). To determine if augmenting mitochondria restores macrophage function, MDMs were co-cultured with MNPs, with or without increasing concentrations of AICAR, and phagocytic ability measured by the uptake of FITC-labeled zymosan. AICAR restored MDM phagocytosis in a dose-dependent manner with a 100µM treatment resulting in significant improvement (Fig.3H).

### MNPs impair antigen presentation and TCR stimulation by macrophages

The ability of macrophages to present processed antigen to stimulate T cells is a key feature of immunosurveillance. To determine how MNPs affect macrophages uptake of tumor cells, Raw264.7 cells were pre-treated overnight with MNPs then co-cultured with CFSE-labeled syngeneic murine 4T1 cancer cells. Pre-treatment with either 1µm or 4µm MNP significantly reduced the amount of tumor cell uptake measured by a reduction in CFSE^+^ expressing macrophages (Fig.4A). We next determined how MNPs impact the ability of macrophages to process and present antigen to stimulate T cell receptor (TCR)-mediated responses. We utilized the T cell hybridoma line DO11.10 which was created by fusion of BALB/c (H-2^d^) T cells specific for chicken ovalbumin (OVA) with the AKR thymoma line BW5147. DO11.10 cells react to OVA323–339 presented in major histocompatibility II (1-A^d^) receptors on Raw264.7 cells^29^. First, we examined the effect of MNPs on TCR stimulation using OVA protein, wherein macrophages must phagocytose, lysosomally process, and present antigen in the presence stimulatory receptors to activate TCRs. In the absence of MNPs, Raw264.7 cells induced OVA-specific stimulation of 44.22% ± 9.03 of co-cultured DO11.10 cells over 72hrs measured by cell proliferation via CSFE dilution (Fig.4B). Addition of 1µm or 4µm MNPs at increasing ratios for 16-18-hrs prior to OVA addition resulted in a significant dose-dependent decrease in macrophage-mediated DO11.10 division (Fig.4B). Next, we cultured Raw264.7 cells with GFP-labeled OVA protein with or without prior MNP exposure and determined OVA uptake via fluorescence microscopy. As expected, control Raw264.7 cells readily phagocytose GFP-OVA (26.4% ± 1.9) and MNP co-culture significantly reduced uptake (14.7% ± 1.2) (Fig.4C).

**Figure 4:**
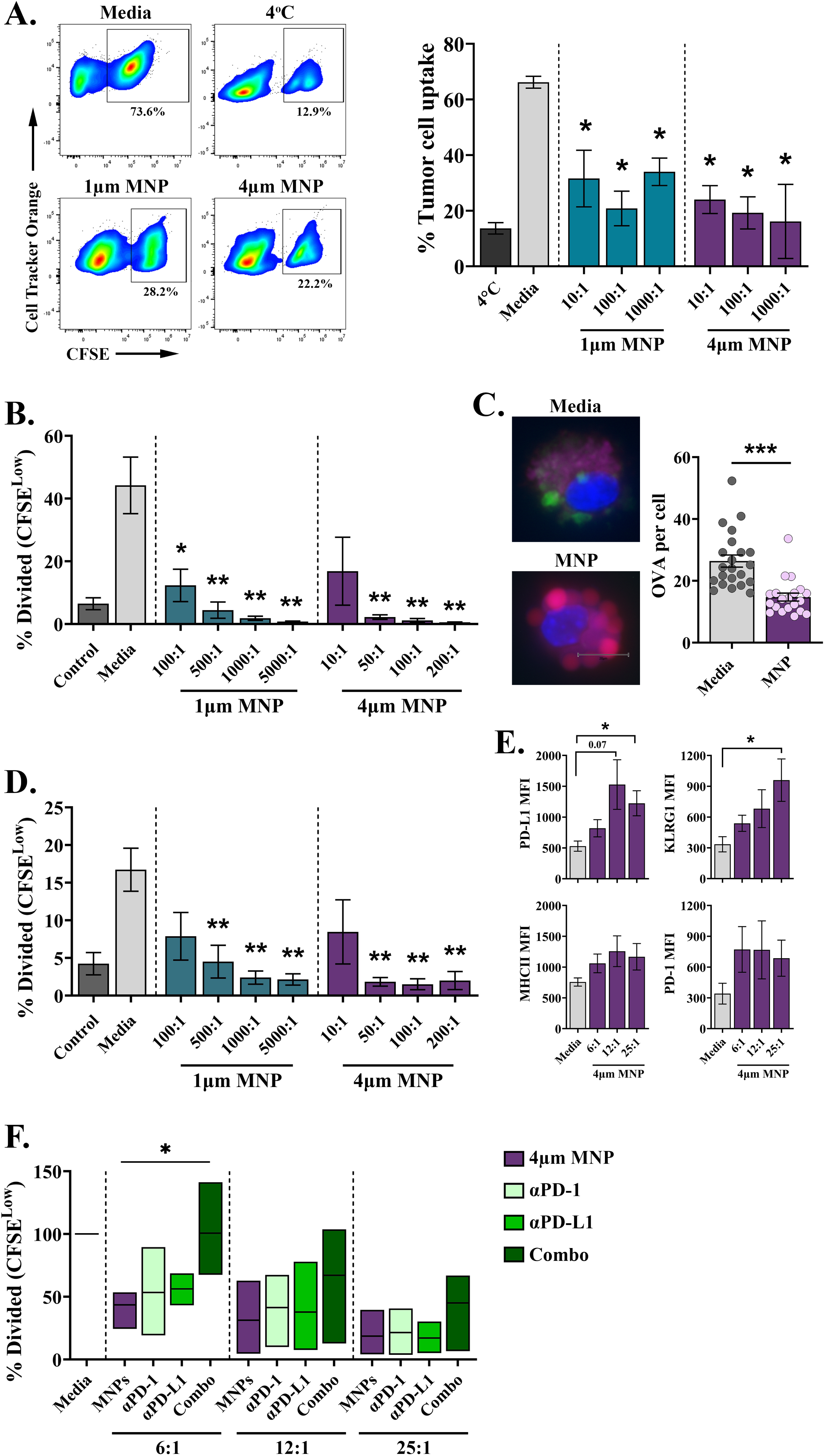
MNPs impair antigen presentation by macrophages. A) Phagocytosis of CSFE-labeled 4T1 tumor cells by Cell Tracker Orange-labeled Raw264.7 cells to (10:1; 4T1s to Raw264.7) after overnight culture MNPs. Media only or 4°C served as positive and negative controls, respectively. Raw264.7 cells cultured with B) OVA protein or D) OVA323–339 peptide and increasing MNP burdens have decreased ability to stimulate OVA-specific TCRs driving DO11.10 cell proliferation. C) Representative images and quantitation of FITC-labeled OVA protein uptake by Raw264.7 cells with or without prior administration of 10:1 4μm MNPs. Nucleus is colored in blue (Hoechst 33342), Ova Protein (green), 4um MPs (red), LAMP1 (purple). E) Mean fluorescence intensity of MHCII, PD-1, PD-L1, and KLRG1 on Raw264.7 cells determined by flow cytometry. F) Additon of blocking antibodies against both PD-1 and PD-L1 restores TCR-mediated DO11.10 activation by MNP treated Raw264.7 cells.

To determine if decreased antigen processing post-MNPs was responsible for the inhibition of DO11.10 stimulation, we pulsed Raw264.7 cells with OVA323–339 peptide which displaces low affinity endogenous peptides within MHCII molecules at the cell surface. As such, peptide pulsed macrophages present antigens without prior phagolysosomal processing. Similar to assays using OVA protein, OVA323–339 peptide pulsed Raw264.7 readily stimulated DO11.10 cells, inducing 22 ± 5.86% division over 72hrs (Fig.4D). Interestingly, prior incubation of Raw264.7 with MNPs again inhibited peptide stimulation of DO11.10 cells (Fig.4D). As this effect was unrelated to endogenous processing, we measure co-stimulatory and co-inhibitory receptor expression on cells post-MNPs. MNPs did not alter levels of extracellular MHCII expression but significantly increased expression of the inhibitory receptors PD-L1, KLRG1, and to a lesser extend PD-1 without effecting the co-stimulatory receptors CD80 and CD86 (Fig.4E and data not shown). To determine if MNP-induced immunosuppression could be reverted, cells were administered MNPs in the presence of blocking antibodies specific for PD-L1 or PD-1. At low MNP to cell ratios, TCR activation by MNP-exposed macrophages could be restored through the addition of both anti-PD-1 and anti-PD-L1 but not with either antibody individually (Fig.4F). Notably, the restoration of TCR activation was lost as macrophages were exposed to greater MNP to cell ratios.

### Sustained presence of MNPs in the lung promotes disease-associated transcriptional pathways in pulmonary macrophages

PM_2.5_ translocates the airway and is associated with systemic pathologies such as cardiovascular disease. To examine the kinetics by which inhaled MNPs disperse through the body, we administered 3x10^7^ fluorescently labeled 1µm nanoplastics (equal numbers of green, red, and crimson) or saline intranasally to FVB/N mice. At 4hrs, 1-day, and 1-week post-administration, the lungs, blood, brains, colons, kidneys, spleens, livers, and hearts were harvested, mechanically dissociated into single-cell suspensions without enzymatic digestion, and immediately analyzed via flow cytometry (Fig.5A). Compared to controls, MNPs were detected to a significant degree in tissue homogenates of the lungs, brains, and colons with a trend towards increased levels in the spleen (Fig.5B). MNPs were transiently observed in the blood, peaking at 4hrs post-administration and returning to baseline by 1-week (Fig.5B). Plastic particles were detected, but not significantly elevated, in all other organs examined and confirmed via fluorescence microscopy (Fig.S3). Strikingly, the amount of MNPs in the lung did not decrease over a 1-week period, highlighting their biopersistence.

**Figure 5:**
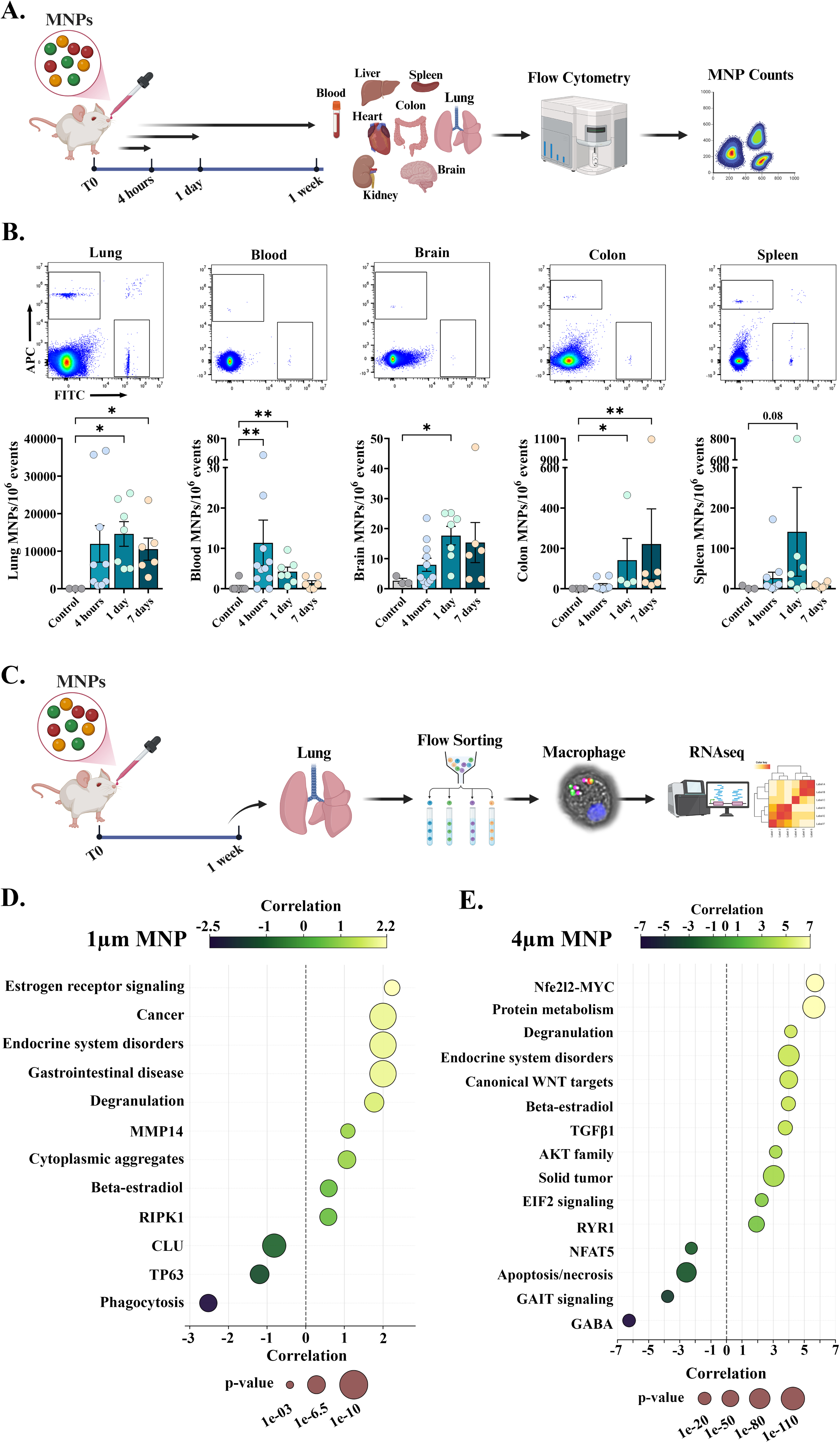
MNPs are retained in the lung and drive aberrant pMac gene expression *in vivo*. A) Schematic of intranasal administration to measure MNP burden in the lungs and extrapulmonary dissemination in mice. B) Quantitation of fluorescently labeled 1um MNPs in the lung, blood, brain, colon, and spleen at 4hrs (n = 9-11), 1-day (n = 4-7), and 1-week (n = 6) or saline treated controls (n = 3-4) with representative dot plots (above). C) Schematic illustrating isolation of pMacs for RNAseq following intranasal administration of 1µm or 4µm MNPs. Macrophage shown is a pMac from the experiment. IPA analysis of gene expression data from isolated pMacs 1-week after intranasal administration of D) 1um (n = 7) or E) 4um MNPs (n = 6) compared to pMacs from mice treated with saline (n = 7).

To determine what transcriptional adaptations pMacs make in response to MNP exposure *in vivo*, CD45^+^ CD11b^+^ CD64^+^ CD88^+^ CD24^−^ Ly-6G^−^ pMacs, including alveolar, interstitial, and recruited macrophages, were isolated via flow sorting from the lungs of mice 1-week after being administered 3x10^7^ 1µm, 5x10^5^ 4µm MNPs, or saline intranasally (Fig.5C). Transcript expression in purified pMacs was measured via bulk RNAseq and analysis performed using Ingenuity Pathway Analysis (IPA). Compared to saline treated controls, pMacs in mice exposed to 1µm or 4µm MNPs had 125 and 549 significant differentially expressed genes (DEG), respectively, with 71 DEGs conserved between MNP sizes. pMacs exposed to 1µm MNPs *in vivo* significantly upregulated pathways of estrogen receptor signaling, degranulation, and cytoplasmic aggregates while downregulating phagocytosis (Fig.5D). These effects were mediated by the upstream regulators MMP14, beta-estradiol, and inhibition of signaling from the extracellular chaperone clusterin (CLU) and TP63, a homolog of the tumor suppressor p53.

Collectively, these responses were associated with shared transcription profiles as those found in cancer, endocrine system disorders, and gastrointestinal disease. Interestingly, pMacs exposed to 4µm MNPs *in vivo* had similar gene expression pathways to those seen after 1µm MNP exposures including degranulation, beta-estradiol signaling, and conserved associations with cancer and endocrine system disorders (Fig.5E) suggesting a central mechanism of MNP-induced pMac dysfunction. Conversely, gene expression in 4µm MNPs exposed pMacs was regulated by unique causal networks including Nfe2l2 (NRF2)-MYC and canonical WNT target genes, increased TGFβ1, AKT family, EIF2, and RYR1 signaling with decreased NFAT5, GAIT, and GABA signaling (Fig.5E). These pMacs had increased protein metabolism and cell survival. Notably, stronger associations between transcript expression and pathology were observed in pMacs from mice administered 4µm MNPs compared to 1µm MNPs, with p-values in cancer-associated transcriptional pathways of 1.06e^-93^ vs. 2.27e^-10^ and for endocrine system disorders of 9.02e^-95^ vs. 2.27e^-10^, respectively.

### MNPs induce unique pathology in human lung tissues and macrophages

To evaluate the direct effect of MNPs on human lung tissues, we treated donor-derived hPCLS with 100µg/ml or 1mg/ml of 1µm MNPs, 4µm MNPs, or DEP for 24hrs and assessed viability via imaging and ATP production. After 24hrs, cells in DEP-treated hPCLS showed no decrease in metabolic activity whereas hPCLS treated with 1µm or 4µm MNPs contained a significant number of dead and/or dying cells determined via confocal microscopy and quantitated green/red LiveDead ratio of composite 100µm z-stacks (Fig.6A). Similarly, the 4µm MNPs exposure, but not the 1µm MNPs significantly reduced ATP production compared to DEP treated hPCLS (Fig.6B). These findings suggest that MNPs induce distinct cellular pathology in human 3D lung cultures compared to DEP and support our previous findings MNPs are cytotoxic at high doses.

**Figure 6:**
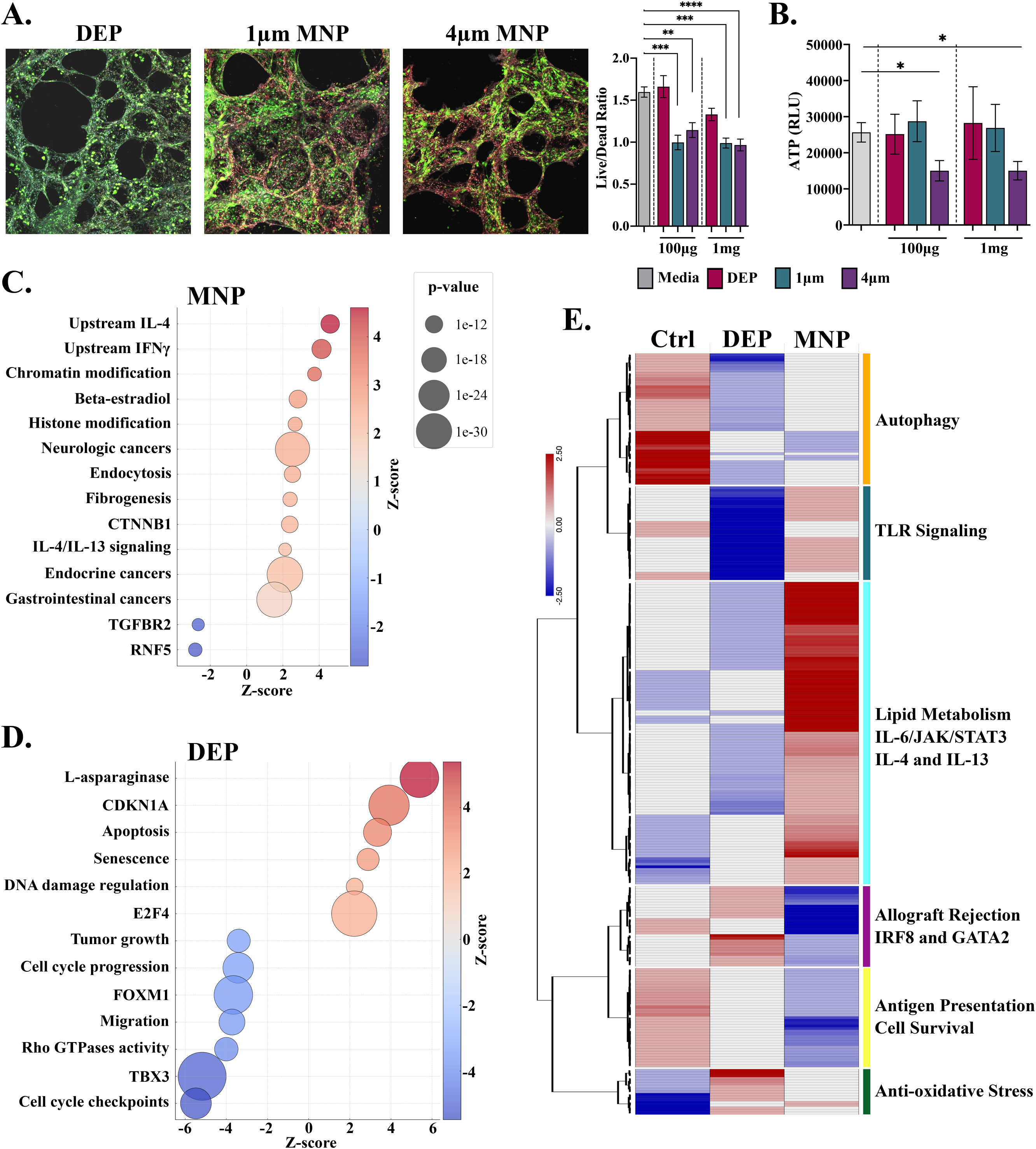
MNPs significantly damage human lung tissues *ex vivo* and alter MDMs *in vitro* compared to DEP. A) Representative images and quantitation of LiveDead viability staining in hPCLS (n = 3 donor tissues). B) ATP measured in culture supernatants of hPCLS treated with 1µm MNPs, 4µm MNPs, or 50ug/ml DEP. IPA analysis of gene expression pathways following treatment of MDMs *in vitro* with C) 1um MNPs (n = 6) or D) DEP (n = 6) compared to saline treated controls (n = 3) for up to 3-days. E) Relative expression and hierarchical clustering of 281 genes which are significantly involved in macrophage regulation and function.

Next, we performed RNAseq on human MDMs treated with 1µm MNPs, DEP, or saline for up to 3 days. Compared to saline treated controls, MNPs upregulated pathways involved in chromatin and histone modification, endocytosis, fibrogenesis, CTNNB1 (β-catenin) and IL-4/IL-13 signaling which were predicted to be downstream of IL-4, IFNγ, and beta-estradiol activation (Fig.6C). Interestingly, the top 36 predicted pathologies associated with MNP-induced gene expression, identified by IPA, were all cancers or related to tumor biology, highlighted by shared genes involved in neurologic, endocrine, and gastrointestinal cancers. Concomitantly, MNP treatment of MDMs inhibited regulation by TGFBR2 and the E3 ubiquitin ligase RNF5, both of which suppress inflammation (Fig.6C). By contrast, DEP exposure upregulated expression of genes regulating l-asparaginase activity, apoptosis, cell senescence, and DNA damage repair, which were associated with inhibition of tumor growth, cell cycle progression, and migration (Fig.6D). These were predicted to be downstream of increased E2F4 and decreased FOXM1 and TBX3 transcription factor signaling and collectively indicative of the cellular response to oxidative stress (Fig.6D). We then examined the relative expression of 281 genes regulating macrophage function from MNP or DEP treated MDMs compared to controls (Table S2). Genes clustered into 6 groups with MNPs driving lipid metabolism, IL-6/JAK/STAT3, and IL-4/IL-13 signaling associated with alternatively activated/wound healing macrophages, while DEP upregulated IRF8 and GATA2 along with the allograft rejection and anti-oxidative stress pathways (Fig.6E). In addition, MNPs inhibited expression of genes involved in antigen presentation in MDMs to a greater extent than DEPs (Fig.6E). Collectively, these findings suggest that MNPs impact diverse biologic functions in macrophages including the endocrine system and β-catenin signaling which may promote epithelial to mesenchymal transition and fibrogenesis.

## Discussion

Since the 1950s, levels of airborne PM2.5 in the United States have steadily declined. Yet, over this same period production of plastics has grown from ∼2 million to over 400 million metric tons each year, an annual growth rate of 8.4%^30^. It is likely that MNPs from production (e.g. tires, textiles, cosmetics, paint, etc.) and fragmentation/degradation represent an increasingly large proportion of the airborne particulate we inhale^31^. Alarmingly, over the past several decades there has been rapid increase in the incidence of lung cancer in never smokers (LCINS) which occurs more commonly in women, at an earlier age, and are less responsive to immune checkpoint inhibitor immunotherapy^32,33^. Similarly, the incidence of COPD in never-smokers has increased, representing ∼24% of adult cases in the U.S. and although plateauing around 2010, asthma incidence has grown to include 7.7–8% of the U.S. population^34–36^. Due to their relatively rapid occurrence, changes in disease prevalence likely reflect the impact of environmental factors, such as MNPs, and indicate that continued research into evolving toxic exposures is necessary.

Following a wave of inflammation, pMacs re-polarize to establish a post-resolution immune state in the lung^37^. Pro-resolution macrophages (Mres) are distinct from the alternatively activated (“M2-like”) pMacs, with high metabolic demands, increased efferocytosis, and production of specialized pro-resolving lipid mediators (SPMs). SPMs including lipoxins, resolvins, protectins, and maresins inhibit neutrophil recruitment and promote epithelial and vascular repair^38–40^. Unlike alternatively activated macrophages which are biased towards fatty-acid oxidation, Mres activity is supported by oxidative phosphorylation via AMPK-mediated autophagy and enhanced mitochondrial biogenesis^41^. Our findings indicate the MNPs are potent inhibitors of mitochondrial function and biogenesis which may inhibit of the ability of Mres in the lung to restore homeostasis. Failure to clear apoptotic bodies results in secondary necrosis, release of DAMPs, and continual recruitment of neutrophils, all of which sustain subclinical “smoldering” inflammation. As such, defects in autophagy have been associated with cancer^42^.

This may be highly relevant to respiratory MNP exposures, as we found that MNPs induced greater cell death and/or metabolic inhibition in human hPCLS *ex vivo* compared to DEP. Alternatively, the inability to shift from alternative activation to Mres may lead to sustained STAT6-IRF4 activation promoting fibrosis. Interestingly, the pro-resolving lipids 15-epi-LXA4 and resolving D1 enhance phagocytosis in Mres through cytoprotective NRF2 (*Nfe2l2*) signaling, an anti-oxidant transcription factor, which was prominently upregulated in pMacs following MNP exposures *in vivo*^43^. Collectively, our findings illustrate that MNPs disrupt multiple processes, including phagolysosomal processing and mitochondrial function, which work in conjunction to maintain lung homeostasis.

As antigen presenting cells and the predominant phagocytes of the immune system, pMacs play an integral role in the induction of adaptive immunity. Through constant sampling of the microenvironment, pMacs are critical for identifying and eliminating malignant progenitor cells via immunosurveillance, a process that occurs continually throughout our lives. Having engulfed malignant cells, activated pMacs traffic to the lymph node where they present tumor-associated antigens to induce tumor-reactive T cell activation^44^. We have found that MNPs affect the antigen presentation functions of macrophages by both restricting phagolysosomal processing as well as inducing inhibitory co-receptors which suppress T cell activation. Loss of antigen presentation will impair immunosurveillance, potentially allowing for tumor immune escape and cancer breakthrough to occur earlier in one’s lifespan. Interestingly, our findings indicate that MNP-impaired phagocytic and antigen presenting function may be partially restored through activating AMPK-mediated biogenesis and blocking PD-1/PD-L1 receptors, respectively. Alternatively, MNPs could impair the antigen processing and presentation of self-antigens on class I MHC molecules from tumors cells directly promoting their escape from immune detection^45^. Although a causal role for MNP-induced immunosuppression in human disease has yet to be established, MNP bioaccumulation and continually impaired immunity are likely to increase susceptibility to chronic lung diseases.

Research into the effects of MNPs is rapidly advancing, yet a number of limitations prevent translation to human health. First, there is no consensus range of MNPs in human lung tissues. Microplastics have been quantified in human lung tissue at ∼0.5 to 1.5 particles per gram which did not include particles under ∼3µm^3^. No single analytic technique can measure particle counts, mass, plastic subtype, size, or shape by itself. Furthermore, there are few standardized methods for sample collection, processing, or handling to avoid contamination. This lack of uniformity precludes comparisons across studies. As such, it is unclear what a relevant dose would be for *in vivo* or *in vitro* modeling. By examining a roughly 5-log10 range of particles to macrophages we show that exposure dose is critical to determining the effects of MNPs. In our studies, lower doses of particles per cell (10^0^-10^2^) failed to induce either inflammation or cell stress/death but dramatically inhibited mitochondria and antigen-specific TCR activation. By contrast, macrophage production of ROS was only detected at cytotoxic levels of MNPs, illustrating the differential effects of MNP dose on biologic response. Notably, with the techniques used in this study and many others, it is impossible to determine if MNPs promote inflammation directly or if the inflammatory response is induced by the release of DAMPs following MNP-mediated cytotoxicity. Second, our study used non-functionalized “pristine” polystyrene microspheres. By contrast, MNPs in the environment are immediately modified gaining eco-corona coatings of biomolecules such as carbohydrates, lipids, proteins and even microorganisms^46,47^. Additionally, although MNPs are generally thought to be inert toxicologically, their ability to bind carcinogens, neurotoxins, and endocrine disrupting chemicals in the environment may have dramatic effects on cells and tissues with significant health implications^48,49^. As such, potential MNP-mediated pathology likely stems from both accumulation of particulate matter in tissues as well as the effects of their endogenous and acquired chemical composition. Future studies must address these limitations to truly understand the pathobiology of MNPs and their contribution to respiratory disease.

## Supporting information

Dhupar et. al., Supplement

---

Supplemental Figure 1: Total particle volume per cell represents a consistent metric for administering MNPs in vitro. 20,000 Raw264.7 cells were co-cultured overnight with 1µm (left) or 4µm MNPs (right) at increasing MNP per cell ratios. Total culture volumes were 200µl, 500µl, and 800µl. ATP production was measured by CellTiter-Glo 2 per manufacturers specifications. No significant change in ATP production was observed.

Supplemental Figure 2: Confocal imaging of MNP-laden monocyte-derived macrophages. Human MDMs treated with MNPs at 10:1 or 100:1 were cultured overnight. Cells were immunostained with Tom20-specific antibody (red), phalloidin (orange), and DAPI (blue). A) Representative illustration of Tom20 expression seen wrapping around intracellular 4µm MNPs.

Supplemental Figure 3: Representative immunofluorescent images showing labeled MNPs in the lung, brain, liver, kidney, colon, heart and spleen. Lung image is of a PCLS generated from mouse tissue 4hrs post-administration of 1µm MNPs (pseudo-color of 3 different fluorescent emissions).

Supplemental Table 1: Details of MNP dosage per experiment reported as particles per cell, concentration (conc.), and particle volume per cell.

Supplemental Tabel 2: List of conventional macrophage-specific transcripts used to compare the effects of MNPs, DEP, and saline following *in vitro* administration to MDMs.

## Author Contributions

Conception and design: RD and ACS. Performing experiments: HMU, NN, CEB, XZ, MC, ST, DS, KM, OAB, LG, AW, RHP, SU, DMG. Analysis and interpretation: RD, MK, SU, DMG, and ACS. Manuscript writing and revision: RD, HMU, CEB, NN, MK, SU, DMG, and ACS. All authors have direct access and have verified all data. All authors have read and approved the final manuscript.

## Acknowledgement

This project utilized the services of the University of Pittsburgh Health Sciences Sequencing Core at UPMC Children’s Hospital of Pittsburgh for RNAseq. Research reported in this publication utilized the bioinformatics, cytometry, and animal facilities of the UPMC Hillman Cancer Center and was supported by the National Cancer Institute of the National Institutes of Health under Award Number P30CA047904.

## Data availability

RNAseq data will be deposited in the Gene Expression Omnibus (GEO) at the time of publication. All additional experimental data is available through the authors upon request.

## Disclaimer

The views expressed in this material are solely those of the authors and do not reflect the official policy or position of the U.S. Government, the Department of Defense, or the Department of Veterans Affairs.

