## Supplementary material for "Microplastics inhibit macrophage bioenergetics impairing homeostatic function and immune responsiveness": Dhupar et. al., Supplement

**Supplemental Data – Extended Methods**

*Polystyrene MNPs and uptake*: Polystyrene microspheres of 1µm, 4µm, and 9µm labeled with either yellow-green (excitation/emission maxima of 505/515 nm), red (580/605 nm), or crimson (625/645 nm) were purchased from ThermoFisher Scientific. Non-fluorescent, unlabeled polystyrene microspheres of 1µm, 4µm, and 9µm were purchased from Spherotech. Human monocyte-derived macrophages (MDM) were generated by culturing magnetic bead-isolated $CD14^+$ cells from donor blood (N = 6, Vitalant) in 20 ng/ml M-CSF for 5-6 days. MDMs were cultured in RPMI-1640 supplemented with 10% heat-inactivated fetal bovine serum (FBS), 2 mM L-glutamine, and 10 U/mL Penicillin-Streptomycin (P/S). $5x10^4$ MDMs were seeded overnight in 4-well Lab-Tek II chamber slides and allowed to adhere at 37°C, 5% $CO_2$ followed by co-culture with MNPs for 24 hours. Cells were then fixed and permeabilized using the True-Nuclear Transcription Factor Buffer Set (BioLegend). Cells were stained with Rab5a antibody (Alexa647; Santa Cruz sc-46692), Rab7 antibody (Alexa647; Santa Cruz, sc-376362), and LAMP-1 antibody (Alexa647; Santa Cruz, sc-19992) in permeabilization buffer overnight. Nuclei were stained with DAPI and the microfilaments were stained with Phalloidin (Alexa488; cell signaling, 8878S). The fluorescence was photographed by an EVOS M5000 microscope. 20,000 Raw264.7 murine macrophages (TIB-71, ATCC) were seeded in 100 µl DMEM supplemented with 10% FBS, 2 mM L-glutamine, and 10 U/mL P/S into 96-well flat-bottom plates and incubated for 4h at 37°C, 5% $CO_2$ to allow for adhesion. Cells were exposed to microplastics at varying concentrations for 24 h. Prior to analysis, cells were washed 3 times to remove any unbound MNPs. MNP uptake was measured via immunofluorescent imaging on an

EVOS M5000 microscope. All conditions were performed in triplicate and experiments were repeated a minimum of three times independently.

*LDH release*: 20,000 Raw264.7 cells (ATCC, TIB-71) were co-cultured with MNPs in a final volume of 100 µl with complete DMEM for 24 hours. Lactate dehydrogenase release was quantified using the CyQUANT™ LDH Cytotoxicity Assay Kit (Invitrogen) according to the manufacturer's instructions. Cell-free supernatants (50µL) were collected in a 96-well plate, mixed with 50µL reaction mixture, and incubated for 30 min at room temperature in the dark. 50ul of stop solution was added and absorbance was read at 490 nm and 650 nm (background) using a Varioskan ALF microplate reader. Percent cytotoxicity was calculated as: (Sample−Spontaneous LDH activity)/(Maximum LDH activity−Spontaneous LDH activity)×100 using background-corrected values ($A_{490} - A_{650}$).

*XTT assay*: For viability, metabolic activity was measured using the CyQUANT XTT Cell Viability Assay Kit (Invitrogen). A working solution (1:6 electron coupling:XTT reagent) was prepared immediately before use, and 70µl was added to each well. After 4h at 37°C, absorbance was recorded at 450 nm and 650 nm using a Varioskan ALF microplate reader. Specific absorbance was calculated as [$Abs_{450}$ nm(Test) – $Abs_{450}$ nm(Blank)] – $Abs_{650}$ nm(Test)], and viability was expressed relative to untreated controls.

*Transcription factor activation:* $2x10^5$ Raw-Dual reporter cells (Raw264.7 dual NF-κB/IRF reporter cells; InvivoGen, rawd-ismip) were seeded in 200 µl in 96-well plates and exposed to increasing concentrations of diesel exhaust particles (25-400 µg/mL), 1µm MNPs (100:1-1600:1,

MNPs/cell), or 4μm MNPs (10:1-160:1, MNPs/cell) with or without 100μg lipopolysaccharide (LPS) for 24hrs. NF-κB activation was measured by the secreted embryonic alkaline phosphatase (SEAP) reporter using QUANTI-Blue detection reagent (InvivoGen) according to the manufacturer's instructions and absorbance measured at 620nm. Interferon response factor (IRF) activation was determined using the Lucia luciferase reporter and quantified using QUANTI-Luc detection reagent (InvivoGen) following the manufacturer's flash luminescence protocol and immediately read on a Varioskan ALF microplate reader. Reporter activity was normalized to untreated cells (LPS- conditions) or LPS-treated controls (LPS+ conditions), and results were expressed as percentage changed relative to the corresponding control.

*JC-1 Assay*: 5x$10^4$ Raw264.7 cells were seeded in Lab-Tek II Chamber Slides and co-cultured in 1ml DMEM with various concentrations of 4μm microplastics for 24-hours at 37°C. CCCP (carbonyl cyanide 3-chlorophenylhydrazone) was used as a positive control (ThermoFisher; 2μM). Cells were then stained with 2μM JC-1 reagent (MitoProbe, ThermoFisher) at 37°C in dark for 30 minutes. Cells were washed thoroughly, labeled with DAPI, and imaged for immunofluorescence. Three-dimensional (3D) fluorescence z-stacks were acquired to characterize mitochondrial activation profiles around the microplastics. Microplastic centroids in (x, y, z) were identified by combining their detection within each 2D optical z-section using a Hough circle transform with their reconciliation in 3D using 3D anisotropic DBSCAN algorithm. For each detected microplastic, radial intensity profiles were extracted in the green and red channels, representing mitochondrial activation and inactivation states, by computing the 3D Euclidean distance from the centroid to every surrounding voxel. Following per-slice background subtraction and per-channel peak normalization, profiles were averaged across all microplastics to characterize the spatial distribution of mitochondrial activation with respect to the microplastics.

*ROS Assay*: $5x10^5$ Raw264.7 cells in 1ml DMEM were co-cultured with 1µm or 4µm MNPs at increasing MNP/cell ratios for 24-hours. Cells were then labeled with 500nM CellROX green kit for 30-60 min at 37°C with 5µm Sytox red included for the final 15 minutes (ThermoFisher). TBHP (Tert-butyl hydroperoxide) (400µm) was added for 1 hour at 37°C as a positive control. JC-1 was immediately measured on a CytoFLEX cytometer and reported as red/green ratio (Beckman Coulter).

*Cytokine Production*: 50µl of fresh cell culture supernatant was analyzed using the Mouse Inflammation Cytometric Bead Array kit simultaneously measuring IFNγ, TNFα, IL-6, IL-10, CCL2, and IL-12p70 (BD Biosciences). Data was collected using a CytoFLEX flow Cytometer and analyzed using FlowJo V10 software (BD Biosciences). Cytokine levels were determined per mean fluorescence intensity measurements from manufacturer provided standards and protocol using GraphPad Prism8 (GraphPad Software).

*Mitochondrial Mass Assay*: Mitochondrial mass was assessed using Tom20 (1:400, CST, 42406) co-stained with Alexa 647 (1:100, Jackson Immuno, 111-605-003). Briefly, cells were fixed in 3.7% PFA in PBS for 5 minutes at 37°C. The cells were permeabilized using 0.25% Triton X-100 in PBS, blocked with 10% goat serum in PBS. Cells were then stained with Rhodamine Phalloidin following manufacturer's protocol (Fisher Scientific, R415) and DAPI. Images were then obtained by confocal microscopy using a 60x objective and Olympus IX83 microscope equipped with a Cicero widefield/confocal system. Analysis was performed using ImageJ and CellProfiler software.

*Seahorse XF Mito Stress Test*: Mitochondrial respiration was measured using the Seahorse XF Mito Stress Test Kit (Agilent Technologies,103010-100) according to the manufacturer's instructions. Briefly, cells were seeded in Seahorse cell culture microplates at a density of 100,000 cells and incubated overnight to allow adherence with or without 100:1 4μm MNPs or 50 μg/ml DEP. Prior to the assay, culture medium was replaced with Seahorse XF Assay Medium (Agilent) supplemented with 10 mM glucose, 2 mM glutamine, and 1 mM pyruvate, and cells were incubated in a non-$CO_2$ incubator at 37°C for 1 hour. Oxygen consumption rate (OCR) was measured under basal conditions and following sequential injections of oligomycin (1 μM), FCCP (1 μM), and rotenone/antimycin A (0.5 μM each). Data were normalized to total cell count per well and analyzed using Wave software (Agilent).

*Phagocytosis*: Raw264.7 cells were co-cultured overnight with 10:1 4μm MNPs then incubated with 5:1 Zymosan A (S. cerevisiae) BioParticles per cell (5:1, ThermoFisher) for 4 hours. Some groups were pre-treated for 1 hours with AICAR (10-100 μM) prior to zymosan addition. Zymosan uptake was measured as the percent of cells expressing AlexaFluor488 by flow cytometry.

*OVA uptake and stimulation assay:* $2.5x10^5$ Raw264.7 cells were seeded in sterile chambered glass slides (Lab-Tek™ II Chamber Slide™ System, ThermoFisher) and co-cultured with 4μm red MNPs (580/605 nm) at 10:1 overnight. Cells were washed three times then cultured with

FITC-labeled ovalbumin protein (25µg/ml; Invitrogen) for 2-4 hours at 37°C. Cells were washed, fixed, permeabilized then stained with LAMP-1 (1:100) at room temperature for 90 minutes followed by extensive washing then DAPI for 5 minutes and visualized via immunofluorescence imaging. $5x10^5$ Raw264.7 cells were seeded in 12-well plates (2ml DMEM) and cultured overnight (16-18hrs) with (100U/ml) IFNγ with or without microplastics. Unbound MNPs were removed, cells washed, then cultured with ovalbumin (OVA) protein (Sigma; 100µg/mL) or $OVA_{323–339}$ peptide (Sigma; 20µg/mL) for 4h at 37°C. DO11.10 T cells (Cat# 85082301, MilliporeSigma) were maintained in culture with complete DMEM at 37°C, 5% $CO_2$. DO11.10 cells were labeled with 0.5µM CFSE (Invitrogen) for 10 minutes at room temperature. CFSE was quenched by adding 4-5 volumes of cold media and incubated on ice for 5 minutes then washed 3 times with complete media. CFSE-labeled DO11.10 cells were added to the OVA protein or peptide treated Raw264.7 cells at a 1:1 ratio in fresh DMEM medium. Co-cultures were maintained at 37°C in 5% $CO_2$ incubator for 72h. Non-adherent DO11.10 cells were then collected, stained for viability with zombie NIR (1:1000; BioLegend) at 4°C in dark for 15 minutes, and immediately analyzed via flow cytometry. Checkpoint inhibitors anti-PD-1 (BioXcell, BP0146) and anti-PD-L1 (BE0101) were included at 30µg/ml in some experiments. Proliferation was normalized to $CSFE^{Low/null}$ cells stimulated in the absence of MNPs. Data was analyzed using FlowJo software (v10.10).

*In vivo exposures*: 10- to 16-week-old female and male FVB/N mice (Jackson Laboratory) were maintained in a specific pathogen-free environment at the University of Pittsburgh animal research facilities at the UPMC Hillman Cancer Center. Mice were administered $3x10^7$ 1µm MNPs equal parts yellow-green, red, and crimson in sterile saline or sterile saline alone intranasally in 40µl administering 20µl/nare. At 4-hours, 1-day, or 1-week post-administration mice were euthanized and lung, liver, heart, kidney, brain, colon, and spleens were harvested and processed into single cell suspension with mechanical digestion only. Red blood cells were lysed from blood samples using BD Pharm Lyse Buffer (Waters Biosciences). Single cell suspensions were immediately analyzed via flow cytometry for detection of 505/515nm, 580/605nm, or 625/645nm fluorescence. Data was analyzed in FlowJo v10.10 and fluorescent MNP counts normalized to $10^6$ total events. Additional mice received non-fluorescent MNPs at $3x10^7$ 1µm or $5x10^5$ 4µm MNPs, or sterile saline intranasally. 1-week post-administration, lungs were excised with thymus and heart carefully removed, finely minced, and digested at 37°C, 5% $CO_2$ for 30 min in the presence of Liberase TM (Millipore Sigma) per manufacturer's instructions. Mechanically and enzymatically dissociated lung homogenate was pressed through a 100µm filter with the plunger of a 5ml syringe followed by extensive washing with cold, sterile PBS. Lung homogenate was then passed through a 70µm filter and stained for CD45, CD11b, CD88, CD24, CD64, Ly-6G, then LiveDead blue viability dye. Pulmonary macrophages were isolated as LiveDead$^{-}$ CD45$^{+}$ CD11b$^{+}$ CD64$^{+}$ CD88$^{+}$ CD24$^{-}$ Ly-6g$^{-}$ cells to ≥96% purity with a Sony900 cell sorter. All experiments were approved by the University of Pittsburgh Institutional Animal Care and Use Committee and conducted in accordance with guidelines of the U.S. Public Health Service Policy on Humane Care and Use of Laboratory Animals.

*Human lung tissue procurement*: The use of human lung tissue was approved by the University of Pittsburgh Institutional Review Board. Lung tissue unsuitable for transplantation was obtained from the Pulmonary, Asthma, Critical Care, and Sleep Medicine Thoracic Tissue Repository and the Center for Organ Recovery and Education. After collection, lung lobes were stored on ice in

DMEM/F12 (Dulbecco's Modified Eagle Medium and Ham's F 12 Nutrient Mixture) or in an organ transplantation preservation solution until processing within 3 hours of receipt. All experimental procedures were performed under sterile conditions.

*Human precision cut lung slice (hPCLS)*: hPCLS were generated as previously described[27]. Briefly, lung lobes were inflated via the main bronchus with 500–600 mL of warm (37–40 °C) 2.5% low–melting point agarose (LMPA; IBI Scientific) prepared in DMEM/F12 with phenol red (ThermoFisher, Waltham, MA) and supplemented with 1% fetal bovine serum (FBS; BioTechne), 1% penicillin/streptomycin, and 1% amphotericin B. The inflated tissue was then kept on ice for 30-45 minutes to allow agarose to solidify. Agarose filled lobes were then cut into approximately 2.5cm thick slices. Tissue cores were obtained from each slice using a 1cm diameter biopsy coring tool (Alabama Research and Development), avoiding major airways and blood vessels. Parenchymal cores were maintained on ice in DMEM/F12 supplemented with 1% FBS, 1% PenStrep, and 1% Amphotericin B (Complete Media 1%). Precision cut lung slices (300 µm thickness) were generated using a Compresstome vibroslicer (VF 510 0Z, Precisionary Instruments) with a cutting speed of 6µm/s and an oscillation frequency of 6Hz. Individual slices were placed into 24 well plates. Plates were incubated overnight at 37 °C in a humidified 5% $CO_2$ atmosphere in complete media 1%. The following day, complete media 1% was replaced with DMEM/F12 supplemented with 0.1% FBS (Complete Media 0.1%) and HPCLS were exposed to MNPs, DEP, or saline for 24hrs.

*hPCLS cell viability assessment*: To evaluate hPCLS viability and survival, two to three 4mm punches were obtained from a single 1cm hPCLS using a biopsy punch. Samples were assessed in triplicate on prior to and at the end of treatments. The CellTiter-Glo 2.0 ATP quantification assay (Promega) was used according to the manufacturers' instructions. LiveDead viability assay was used including cell-permeant Calcein AM which is converted to green-fluorescent calcein by intracellular esterases indicating metabolically active cells, and the cell-impermeant BOBO-3 Iodide which produces red fluorescence when bound to DNA/nucleic acids inside dead or dying cells.

*RNA-sequencing*: Murine pMacs were isolated 1 week after MNP administration. Human MDMs were treated in vitro with 4μm MNPs at 100:1, 50μg DEP, or saline for up to 3 days. RNA was extracted using the RNeasy kit (Qiagen). Generation of cDNA, library preparation, and sequencing were performed by the Health Science Sequencing Core at UPMC Children's Hospital and analysis provided by the Biostatistics Facility of the UPMC Hillman Cancer Center.

*Statistical analysis:* Data between groups were analyzed using non-parametric Kruskal-Wallis test followed by Bonferroni-adjusted p-values for post-hoc comparisons using and GraphPad Prism11 (GraphPad Software). All in vitro assays were performed a minimum of 3 times independently. For all hypothesis, statistical significance was denoted as $p \leq 0.05$ (*), $p \leq 0.01$ (**), $p \leq 0.001$ (***), and $p \leq 0.0001$ (****). All data are reported as the mean ± SEM.

| Figure | Size | Ratio | Conc. (µg/ml) | PV (µm$^3$) | PV/cell (µm$^3$) | Cells/vol |
|---|---|---|---|---|---|---|
| **1D/1E** | 1µm | 1,000:1 | 110 | 1.048E+07 | 5.240E+02 | 2x10$^4$/100 |
| | 1µm | 5,000:1 | 550 | 5.240E+07 | 2.620E+03 | |
| | 1µm | 10,000:1 | 1,100 | 1.048E+08 | 5.240E+03 | |
| | 1µm | 50,000:1 | 5,500 | 5.240E+08 | 2.620E+04 | |
| | 4µm | 100:1 | 704 | 1.048E+06 | 5.240E+01 | |
| | 4µm | 500:1 | 3,520 | 5.240E+06 | 2.620E+02 | |
| | 4µm | 1,000:1 | 7,040 | 6.702E+08 | 3.351E+04 | |
| | 4µm | 5,000:1 | 35,200 | 3.351E+09 | 1.676E+05 | |
| **2A/2B** | 1µm | 100:1 | 55 | 1.048E+07 | 5.240E+01 | 2x10$^5$/200 |
| | 1µm | 200:1 | 110 | 2.096E+07 | 1.048E+02 | |
| | 1µm | 400:1 | 220 | 4.192E+07 | 2.096E+02 | |
| | 1µm | 800:1 | 440 | 8.384E+07 | 4.192E+02 | |
| | 1µm | 1,600:1 | 880 | 1.677E+08 | 8.384E+02 | |
| | 4µm | 10:1 | 352 | 6.702E+07 | 3.351E+02 | |
| | 4µm | 20:1 | 704 | 1.340E+08 | 6.702E+02 | |
| | 4µm | 40:1 | 1,408 | 2.681E+08 | 1.340E+03 | |
| | 4µm | 80:1 | 2,816 | 5.362E+08 | 2.681E+03 | |
| | 4µm | 160:1 | 5,632 | 1.072E+09 | 5.362E+03 | |
| **2C/2D** | 1µm | 10:1 | 1.1 | 1.048E+06 | 5.240E+00 | 2x10$^5$/1,000 |
| | 1µm | 100:1 | 11 | 1.048E+07 | 5.240E+01 | |
| | 1µm | 500:1 | 55 | 5.240E+07 | 2.620E+02 | |
| | 1µm | 1,000:1 | 110 | 1.048E+08 | 5.240E+02 | |
| | 1µm | 5,000:1 | 550 | 5.240E+08 | 2.620E+03 | |
| | 4µm | 10:1 | 70.4 | 6.702E+07 | 3.351E+02 | |
| | 4µm | 50:1 | 352 | 3.351E+08 | 1.676E+03 | |
| | 4µm | 100:1 | 704 | 6.702E+08 | 3.351E+03 | |
| | 4µm | 200:1 | 1,408 | 1.340E+09 | 6.702E+03 | |
| | 4µm | 500:1 | 3,520 | 3.351E+09 | 1.676E+04 | |
| **3A/3B** | 4µm | 100:1 | 1,760 | 1.676E+08 | 3.351E+03 | 5x10$^4$/100 |
| **3C** | 4µm | 1:1 | 17.6 | 1.676E+07 | 3.351E+01 | |
| | 4µm | 3:1 | 52.8 | 5.027E+07 | 1.005E+02 | |
| | 4µm | 6:1 | 105 | 1.005E+08 | 2.011E+02 | |
| | 4µm | 12:1 | 211 | 2.011E+08 | 4.021E+02 | |
| | 4µm | 25:1 | 440 | 4.189E+08 | 8.378E+02 | |
| **3E/6C/6E** | 1µm | 10:1 | 6.88 | 1.310E+06 | 5.240E+00 | 2.5x10$^5$/200 |
| **3F** | 1µm | 100:1 | 6.88 | 1.310E+06 | 5.240E+01 | 2.5x10$^4$/200 |
| | 4µm | 100:1 | 440 | 8.378E+07 | 3.351E+03 | |
| **3H** | 1µm | 100:1 | 110 | 2.620E+06 | 5.240E+00 | 5x10$^5$/250 |
| **4A** | 1µm | 10:1 | 0.55 | 5.240E+05 | 5.240E+00 | 1.0x10$^5$/1000 |
| | 1µm | 100:1 | 5.5 | 5.240E+06 | 5.240E+01 | |
| | 1µm | 1,000:1 | 55 | 5.240E+07 | 5.240E+02 | |
| | 4µm | 10:1 | 35.2 | 3.351E+07 | 3.351E+02 | |
| | 4µm | 100:1 | 352 | 3.351E+08 | 3.351E+03 | |
| | 4µm | 1,000:1 | 3520 | 3.351E+09 | 3.351E+04 | |
| **4B/4D** | 1µm | 100:1 | 0.69 | 1.310E+07 | 5.240E+01 | 2.5x10$^5$/2,000 |
| | 1µm | 500:1 | 3.44 | 6.550E+07 | 2.620E+02 | |
| | 1µm | 1,000:1 | 6.88 | 1.310E+08 | 5.240E+02 | |
| | 1µm | 5,000:1 | 34.38 | 6.550E+08 | 2.620E+03 | |
| | 4µm | 10:1 | 44 | 8.378E+07 | 3.351E+02 | |
| | 4µm | 50:1 | 220 | 4.189E+08 | 1.676E+03 | |
| | 4µm | 100:1 | 440 | 8.378E+08 | 3.351E+03 | |
| | 4µm | 200:1 | 880 | 1.676E+09 | 6.702E+03 | |
| **4F** | 4µm | 6:1 | 26.4 | 5.027E+07 | 2.011E+02 | 2.5x10$^5$/2,000 |
| | 4µm | 12:1 | 52.8 | 1.005E+08 | 4.021E+02 | |
| | 4µm | 25:1 | 110 | 2.094E+08 | 8.378E+02 | |

| Gene | DEP vs Ctrl | MNP vs Ctrl | MNP vs DEP |
|---|---|---|---|
| ABCA1 | 0.126513969 | 0.781459382 | 0.098276756 |
| ABCG1 | 0.487107251 | 0.275943972 | 0.018439725 |
| ACTG1 | 0.728114292 | 0.149281147 | 0.017915666 |
| ACTN1 | 0.501872477 | 0.006533714 | 0.006761141 |
| ADAM10 | 0.492941844 | 0.739081537 | 0.641580735 |
| AIF1 | 0.62067426 | 0.313984622 | 0.499142003 |
| ALOX5 | 0.030165384 | 0.395212671 | 0.082482657 |
| APOE | 0.445301144 | 0.015611133 | 0.02865042 |
| ARF3 | 0.380455397 | 0.541400832 | 0.050719326 |
| ATF3 | 0.954971095 | 0.100383312 | 0.035846228 |
| ATF4 | 0.013750559 | 0.008515036 | 0.831826387 |
| ATG10 | 0.61309691 | 0.922474935 | 0.435414314 |
| ATG101 | 0.853982057 | 0.99809598 | 0.811312871 |
| ATG12 | 0.90682091 | 0.282538174 | 0.214305885 |
| ATG13 | 0.697699362 | 0.419085875 | 0.58704987 |
| ATG14 | 0.565049181 | 0.606124273 | 0.153794021 |
| ATG16L1 | 0.865131352 | 0.85081797 | 0.642590381 |
| ATG3 | 0.036593721 | 0.059881416 | 0.777300351 |
| ATG5 | 0.281956047 | 0.048678155 | 0.250991716 |
| ATG7 | 0.882936944 | 0.426621002 | 0.392945207 |
| ATG9A | 0.072864047 | 0.062204372 | 0.929651069 |
| ATP5F1B | 0.97666377 | 0.996881301 | 0.659078548 |
| ATP5PB | 0.572180605 | 0.425619435 | 0.759541554 |
| ATP6V0B | 0.623456025 | 0.186605374 | 0.272708392 |
| ATP6V0C | 0.079138039 | 0.012594839 | 0.328645357 |
| ATP6V0D1 | 0.659436257 | 0.49425674 | 0.137988067 |
| ATP6V0D2 | 0.05784168 | 0.824445466 | 0.026980256 |
| ATP6V0E1 | 0.751434506 | 0.006885936 | 0.001641562 |
| ATP6V1A | 0.594490054 | 0.269262514 | 0.449284423 |
| B2M | 0.64424122 | 0.000124965 | 8.08455E-06 |
| BAX | 0.434039055 | 0.765049308 | 0.526407169 |
| BCL2 | 0.87925196 | 0.311873132 | 0.262338243 |
| BCL2A1 | 0.918220069 | 0.438684683 | 0.246063721 |
| BECN1 | 0.835122276 | 0.342416978 | 0.33605147 |

| | | | |
|---|---|---|---|
| BNIP3 | 0.790742009 | 0.099804768 | 0.070590316 |
| BNIP3L | 0.881285998 | 0.166299159 | 0.042892304 |
| CASP1 | 0.823695364 | 0.661699355 | 0.384256943 |
| CCL18 | 0.748765434 | 0.026271118 | 0.013280744 |
| CCL2 | 0.779888431 | 0.923941825 | 0.619960328 |
| CCL22 | 0.581829018 | 0.006678362 | 0.004238674 |
| CCL23 | 0.523276797 | 0.218075153 | 0.015920876 |
| CCL3 | 0.939019317 | 0.042147102 | 0.005418573 |
| CCL3L1 | 0.075513078 | 0.017785541 | 7.82768E-08 |
| CCL4 | 0.745699971 | 0.185482843 | 0.031719556 |
| CCL4L1 | 0.167776449 | 0.68568544 | 0.019552472 |
| CCL5 | 0.822550536 | 0.278137249 | 0.085553645 |
| CCR1 | 0.746006531 | 0.501477038 | 0.188344114 |
| CCR5 | 0.227550041 | 0.657045281 | 0.030051366 |
| CCRL2 | 0.610353293 | 0.78189333 | 0.299138268 |
| CD14 | 0.728273034 | 0.850662227 | 0.478585363 |
| CD151 | 0.282569331 | 0.154733665 | 0.648746202 |
| CD200R1 | 0.349990163 | 0.432957191 | 0.842014548 |
| CD209 | 0.846189808 | 0.905431112 | 0.679889878 |
| CD22 | 0.330262067 | 0.603358657 | 0.550007103 |
| CD274 | 0.072299663 | 0.000966968 | 0.048700705 |
| CD276 | 0.609011733 | 0.052493648 | 0.059020278 |
| CD28 | 0.666852327 | 0.549284382 | 0.824543424 |
| CD300A | 0.937938322 | 0.020315418 | 0.003144302 |
| CD300C | 0.777312079 | 0.249572951 | 0.252365113 |
| CD300LB | 0.489667208 | 0.093619521 | 0.192850161 |
| CD300LF | 0.556646861 | 0.674877363 | 0.823865558 |
| CD302 | 0.717107907 | 0.500731807 | 0.681732403 |
| CD33 | 0.867434244 | 0.718698628 | 0.494356897 |
| CD36 | 0.986532104 | 0.783137192 | 0.699852752 |
| CD37 | 0.457530297 | 0.801839931 | 0.189304897 |
| CD40 | 0.796936623 | 0.413877583 | 0.460783322 |
| CD44 | 0.313369352 | 0.107090339 | 0.425252204 |
| CD47 | 0.537042449 | 0.482601436 | 0.911574627 |
| CD48 | 0.538141509 | 0.12361005 | 0.223090882 |

| | | | |
|---|---|---|---|
| CD52 | 0.971776939 | 0.371750584 | 0.256476409 |
| CD63 | 0.723305767 | 0.017540436 | 0.007523927 |
| CD68 | 0.071613278 | 0.881621273 | 0.009887785 |
| CD74 | 0.077853911 | 0.072415553 | 0.965040085 |
| CD80 | 0.987720326 | 0.630353049 | 0.517809834 |
| CD81 | 0.371478624 | 0.052684313 | 0.167826541 |
| CD82 | 0.499575989 | 0.509258376 | 0.078838863 |
| CD83 | 0.740926997 | 0.142781856 | 0.133699232 |
| CD84 | 0.048054046 | 0.01805406 | 0.609023853 |
| CD86 | 0.583064206 | 0.302944345 | 0.038053085 |
| CD9 | 0.497915409 | 0.008004645 | 1.10342E-05 |
| CD93 | 0.777630723 | 0.637056513 | 0.802452613 |
| CD99 | 0.864493563 | 0.159816389 | 0.03724941 |
| CD99L2 | 0.831814905 | 0.002093333 | 0.000197765 |
| CDKN1A | 0.696366743 | 0.488693315 | 0.689711797 |
| CEBPB | 0.372526739 | 0.064129821 | 0.205170016 |
| CHI3L1 | 0.80580144 | 0.055087811 | 0.026946102 |
| CHIT1 | 0.67192636 | 0.059574459 | 0.0533301 |
| CHMP2A | 0.939086234 | 0.929652788 | 0.828761677 |
| CHMP4B | 0.047672001 | 0.025257854 | 0.743435328 |
| CIITA | 0.342094794 | 0.218628421 | 0.713230681 |
| CISH | 0.811550341 | 0.000822292 | 3.5338E-06 |
| CLEC4E | 0.756788564 | 0.65459305 | 0.857031993 |
| COL1A1 | 0.711914623 | 0.045711703 | 0.029841859 |
| COX1 | 0.764817027 | 0.423539867 | 0.50741808 |
| COX2 | 0.795200348 | 0.733165486 | 0.914343412 |
| COX3 | 0.826536571 | 0.98986873 | 0.759111716 |
| COX4I1 | 0.673903106 | 0.215345969 | 0.279833293 |
| CSF1R | 0.872617377 | 0.21148877 | 0.150262652 |
| CTSB | 0.571028613 | 0.814415674 | 0.289158288 |
| CTSD | 0.312322092 | 0.82243155 | 0.10237636 |
| CTSL | 0.77149409 | 0.803244843 | 0.956451897 |
| CXCL16 | 0.777155721 | 0.29799706 | 0.317078323 |
| CXCL2 | 0.784656154 | 0.014408693 | 0.006756789 |
| CXCL3 | 0.264966203 | 0.574396776 | 0.031532729 |

| | | | |
|---|---|---|---|
| CXCL5 | 0.612868017 | 0.190670875 | 0.016654668 |
| CXCL8 | 0.006788892 | 0.332120738 | 1.38049E-06 |
| CXCR2 | 0.709149552 | 0.62716764 | 0.262521295 |
| CXCR4 | 0.683006293 | 0.039706761 | 0.029540643 |
| CXCR5 | 0.004136043 | 0.000423297 | 0.304607018 |
| CYBB | 0.493332077 | 0.765737724 | 0.193514848 |
| DDIT3 | 0.583639983 | 0.07118973 | 0.002134438 |
| EIF2AK3 | 0.606234139 | 0.940199127 | 0.56183459 |
| FCGR1A | 0.76285316 | 0.853142831 | 0.877560638 |
| FCGR3A | 0.304988171 | 0.942566116 | 0.207342983 |
| FN1 | 0.856873915 | 0.005183824 | 0.000543415 |
| FUNDC1 | 0.855468696 | 0.764343027 | 0.536309303 |
| GAA | 0.259570547 | 0.779683899 | 0.262447752 |
| GADD45A | 0.494447233 | 0.69183472 | 0.70412228 |
| GBA | 0.037263882 | 0.104543363 | 0.542916647 |
| GCLC | 0.94691888 | 0.105101116 | 0.039857704 |
| GCLM | 0.199664599 | 0.429232919 | 0.516169039 |
| GLUL | 0.447959901 | 0.870393426 | 0.43077964 |
| GPX1 | 0.547976259 | 0.002463096 | 0.001335869 |
| GPX4 | 0.982600677 | 0.157989537 | 0.066542419 |
| HLA-A | 0.6950065 | 0.012042114 | 0.000124282 |
| HLA-B | 0.761183894 | 0.044256164 | 0.024018196 |
| HLA-C | 0.819944162 | 0.220233635 | 0.054735581 |
| HLA-DPA1 | 0.638079121 | 0.477441163 | 0.118604993 |
| HLA-DPB1 | 0.84663117 | 0.175806564 | 0.125307328 |
| HLA-DRA | 0.894155601 | 0.130779218 | 0.068391215 |
| HLA-DRB1 | 0.887860338 | 0.059388141 | 0.02110789 |
| HMGB1 | 0.004454434 | 0.00774248 | 0.808990784 |
| HMOX1 | 0.700171308 | 0.755944634 | 0.921729279 |
| HSPA1A | 0.784498465 | 0.602844775 | 0.041927621 |
| HSPA1B | 0.960817331 | 0.58031348 | 0.138519487 |
| HSPA5 | 0.27240316 | 0.026278243 | 0.137012021 |
| ICAM2 | 0.810301854 | 0.96572152 | 0.797002005 |
| IFIT1 | 0.094488376 | 0.097296462 | 0.98003919 |
| IFIT2 | 0.697770183 | 0.664548813 | 0.28018526 |

| | | | |
|---|---|---|---|
| IFIT3 | 0.224631986 | 0.374553769 | 0.661622764 |
| IFNGR1 | 0.423861736 | 0.576107804 | 0.750282756 |
| IKBKB | 0.199506171 | 0.259081111 | 0.836673307 |
| IL10 | 0.735840562 | 0.603373479 | 0.262130046 |
| IL10RA | 0.781189891 | 0.327034159 | 0.096706416 |
| IL16 | 0.504588888 | 0.808581693 | 0.229798288 |
| IL18 | 0.045506726 | 0.57101991 | 0.059882193 |
| IL18BP | 1.85055E-06 | 0.000922888 | 0.054026042 |
| IL1B | 0.050122388 | 0.490834478 | 0.000559684 |
| IL1R1 | 0.976045932 | 0.10207319 | 0.036381492 |
| IL21R | 0.842256783 | 0.025204143 | 0.001357402 |
| IL27RA | 0.503922217 | 0.094946794 | 0.185728976 |
| IL2RG | 0.730756043 | 0.149566756 | 0.1467696 |
| IL4I1 | 0.455358927 | 0.048192556 | 0.105041252 |
| IL4R | 0.004462735 | 0.001231739 | 0.626711815 |
| IL6R | 0.904949185 | 0.490501167 | 0.453120348 |
| IL7 | 0.792097828 | 0.167975871 | 0.157189278 |
| IL7R | 0.974913926 | 0.001139279 | 2.12731E-05 |
| IRAK1 | 0.8514346 | 0.289946552 | 0.250714999 |
| IRAK3 | 0.93072102 | 0.778679499 | 0.798903937 |
| IRAK4 | 0.964333934 | 0.344047744 | 0.243742609 |
| IRF1 | 0.059403698 | 0.330030928 | 0.229921185 |
| IRF2 | 0.661471842 | 0.12039468 | 0.008781977 |
| IRF3 | 0.265248693 | 0.973646102 | 0.155685901 |
| IRF5 | 0.010660512 | 0.025230925 | 3.43416E-05 |
| IRF7 | 0.316183884 | 0.553571916 | 0.997545967 |
| IRF8 | 0.655017569 | 0.397899953 | 0.154747317 |
| IRF9 | 0.707136813 | 0.313213573 | 0.564235605 |
| ITGAM | 0.570866272 | 0.528078089 | 0.043015647 |
| ITGB2 | 0.031486965 | 0.949339868 | 0.068427095 |
| JUNB | 0.883340839 | 0.123701921 | 0.065596772 |
| LAMP1 | 0.177340952 | 0.67347145 | 0.219885802 |
| LAMP2 | 0.767288672 | 0.023617663 | 0.000730363 |
| LDLR | 0.405205633 | 0.742991048 | 0.125549519 |
| LGALS3 | 0.002144686 | 0.000158387 | 1.44618E-19 |

| | | | |
|---|---|---|---|
| LIPA | 0.514727046 | 0.901670047 | 0.484947756 |
| LRP1 | 0.176921101 | 0.111491851 | 0.749678429 |
| LYZ | 0.763459661 | 0.574858766 | 0.730925223 |
| MACROH2A1 | 0.154047616 | 0.20535281 | 0.000379426 |
| MAFB | 0.983155664 | 0.882855193 | 0.867397719 |
| MAP1LC3A | 0.950097828 | 0.339852558 | 0.189954915 |
| MAP1LC3B | 0.002885306 | 0.931879555 | 5.72968E-05 |
| MAP1LC3B2 | 0.990518616 | 0.614233003 | 0.614233003 |
| MAP1LC3C | 0.942570305 | 0.734710411 | 0.727567729 |
| MAPK1 | 0.290237001 | 0.517545026 | 0.024969375 |
| MARCO | 0.460186143 | 0.536521929 | 0.873263525 |
| MCOLN1 | 0.789000988 | 0.481342526 | 0.199442734 |
| MERTK | 0.887525371 | 0.829943589 | 0.638447281 |
| MMP14 | 2.20673E-06 | 1.35998E-07 | 0.480040355 |
| MMP19 | 0.16829196 | 0.075189657 | 0.596104408 |
| MMP2 | 0.034671338 | 0.174442275 | 0.318488493 |
| MMP24OS | 0.30571267 | 0.011063631 | 0.049270604 |
| MMP25 | 1.2759E-05 | 0.000242254 | 0.346668344 |
| MMP7 | 0.908192982 | 0.280591311 | 0.114124786 |
| MMP8 | 0.498521458 | 0.127136836 | 0.003811837 |
| MMP9 | 0.098235921 | 0.00517452 | 0.130693428 |
| MRC1 (CD206) | 0.934010058 | 0.048120369 | 0.012305 |
| MSR1 | 0.421448298 | 0.038899821 | 0.095340688 |
| MYD88 | 0.886475703 | 0.534096895 | 0.312099716 |
| ND1 | 0.571619884 | 0.063374138 | 0.087928688 |
| ND2 | 0.744483557 | 0.058646972 | 0.038628273 |
| ND3 | 0.274425018 | 3.35107E-06 | 2.61483E-06 |
| ND4 | 0.758578885 | 0.090171816 | 0.066576584 |
| ND4L | 0.582210577 | 0.855850588 | 0.626432215 |
| ND5 | 0.245741261 | 0.023685536 | 0.145337742 |
| ND6 | 0.270979798 | 0.265685866 | 0.987960577 |
| NFKB1 | 0.501209741 | 0.034730965 | 0.059078016 |
| NFKB2 | 0.052188505 | 6.26406E-06 | 0.000689133 |
| NFKBIA | 0.732264215 | 0.897256404 | 0.536288398 |
| NLRP3 | 0.676311929 | 0.606304458 | 0.898149381 |

| | | | |
|---|---|---|---|
| NOTCH2 | 0.052287808 | 0.545577239 | 0.077295066 |
| NQO1 | 7.01785E-06 | 0.515207435 | 4.15879E-07 |
| NRP1 | 0.919188181 | 0.6044145 | 0.581830884 |
| NRP2 | 0.058046097 | 0.011001364 | 0.393117532 |
| OAS1 | 0.887217964 | 0.225456006 | 0.159338306 |
| OAS2 | 0.714360657 | 0.808668084 | 0.422607717 |
| OAS3 | 0.913679713 | 0.813422219 | 0.649443588 |
| PARP1 | 0.55038615 | 0.070912805 | 0.001556979 |
| PIK3AP1 | 0.83329915 | 0.464067532 | 0.491567084 |
| PLIN2 | 0.837028762 | 0.02666681 | 7.80614E-06 |
| PPARA | 0.200977489 | 0.571002936 | 0.016138784 |
| PPARD | 0.291440051 | 4.69611E-05 | 6.89129E-05 |
| PPARG | 0.874874758 | 0.735696618 | 0.81269537 |
| PYCARD | 0.103089317 | 0.216124983 | 0.602329819 |
| RAB7A | 0.033786923 | 0.503877195 | 0.000236961 |
| RAB7B | 0.907691398 | 0.191314153 | 0.060134798 |
| RAC1 | 0.753946035 | 0.236837694 | 0.251104093 |
| RAD51 | 0.000111888 | 0.933831465 | 7.27379E-07 |
| RB1CC1 | 0.494634797 | 0.69446791 | 0.16057598 |
| RHOA | 0.568202358 | 0.039326093 | 0.049196057 |
| S100A11 | 0.205874132 | 0.589956094 | 0.337276715 |
| S100A6 | 0.56190613 | 0.001903322 | 0.000857487 |
| S100A8 | 0.834810671 | 0.130192366 | 0.084882605 |
| S100A9 | 0.395151403 | 0.809051732 | 0.421320654 |
| SERPINA1 | 0.123607074 | 0.106788376 | 3.23969E-05 |
| SERPINB1 | 0.953395945 | 0.149837839 | 0.049422242 |
| SERPINB2 | 0.174438526 | 0.542118791 | 0.009314708 |
| SERPINB6 | 0.191691607 | 0.104141946 | 0.000121621 |
| SERPINB8 | 0.820309954 | 0.944862998 | 0.835578987 |
| SERPINB9 | 0.631951382 | 0.01033896 | 0.006355434 |
| SIRPA | 0.007115226 | 0.004887913 | 0.871917133 |
| SLAMF8 | 0.014418923 | 0.011702288 | 0.923756399 |
| SNAP29 | 0.757230202 | 0.513235747 | 0.207393256 |
| SOCS3 | 0.509401438 | 0.219020017 | 0.464842031 |
| SOD1 | 0.602996268 | 0.115093933 | 0.166931636 |

| | | | |
|---|---|---|---|
| SOD2 | 0.322736497 | 0.901004095 | 0.253048371 |
| SPARC | 0.896313073 | 0.11060877 | 0.052600875 |
| SPP1 (OPN) | 0.669750005 | 0.417299193 | 0.101589748 |
| SQSTM1 | 0.00036306 | 0.517014298 | 0.000115797 |
| SREBF1 | 0.585165994 | 0.908467726 | 0.569009673 |
| SREBF2 | 0.13862909 | 0.620586479 | 0.193163409 |
| STAT1 | 0.585785014 | 0.208761837 | 0.348644076 |
| STAT2 | 0.043186479 | 0.070129446 | 0.778108022 |
| STAT3 | 0.338604819 | 0.339832559 | 0.996519183 |
| STAT4 | 0.52378025 | 0.92346077 | 0.470651869 |
| STAT5A | 0.173653822 | 0.757116635 | 0.027827225 |
| STAT5B | 0.454018225 | 0.262224883 | 0.014630573 |
| STAT6 | 0.180648632 | 0.400083857 | 0.51137366 |
| STX17 | 0.586405513 | 0.166206978 | 0.012633647 |
| TFAM | 0.271249294 | 0.810717675 | 0.079382981 |
| TGFBI | 0.035867512 | 0.214620255 | 0.008678165 |
| TLR1 | 0.186412313 | 0.840986356 | 0.046261564 |
| TLR2 | 0.931670985 | 0.37638321 | 0.200035179 |
| TLR3 | 0.856352036 | 0.705837267 | 0.797134841 |
| TLR4 | 0.871526315 | 0.412024486 | 0.383935312 |
| TLR5 | 0.97371054 | 0.480818342 | 0.376934108 |
| TLR6 | 0.857653976 | 0.817134732 | 0.946731503 |
| TLR7 | 0.540939733 | 0.782638805 | 0.657365968 |
| TLR8 | 0.025002985 | 0.323640948 | 0.098609139 |
| TNF | 0.289206993 | 0.360547823 | 0.841271671 |
| TP53 | 0.250773909 | 0.00032696 | 0.001449258 |
| TRAF6 | 0.196026168 | 0.600682167 | 0.312517702 |
| TREM2 | 0.925281645 | 0.897597435 | 0.963182738 |
| TXNRD1 | 0.022336705 | 0.123894768 | 0.323738048 |
| ULK1 | 0.129373731 | 0.127612206 | 0.995415621 |
| ULK2 | 0.379363608 | 0.30101526 | 0.012095385 |
| VAMP8 | 0.459902144 | 0.104390381 | 0.24346044 |
| VPS4A | 0.33787987 | 0.780984539 | 0.104869964 |
| VPS4B | 0.825658163 | 0.518221854 | 0.255870825 |
| WDR1 | 0.926507347 | 0.818226652 | 0.670301426 |

| | | | |
|---|---|---|---|
| XBP1 | 0.590358349 | 0.214148738 | 0.355823769 |
| XRCC1 | 0.532232255 | 0.385223327 | 0.052718515 |

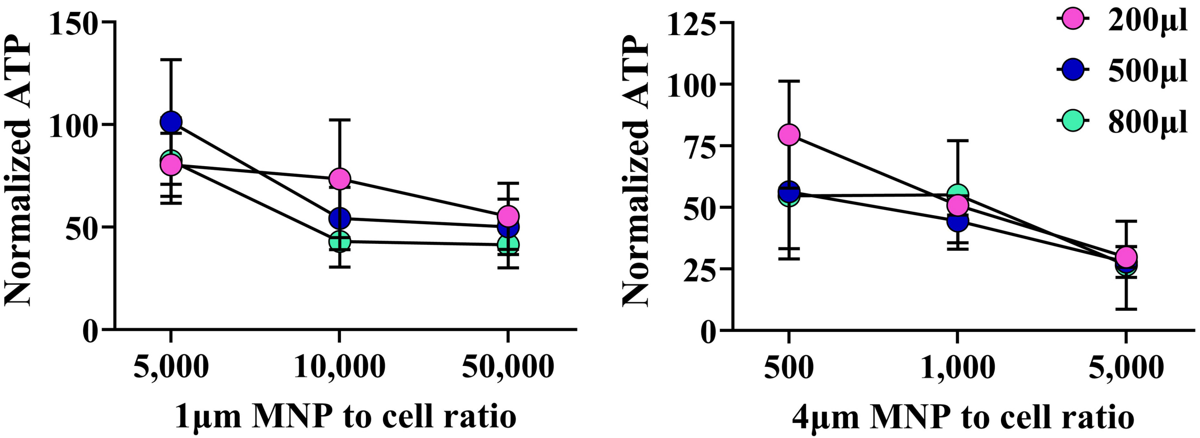

Normalized ATP
150
100
50
0
5,000
10,000
50,000
1µm MNP to cell ratio
Normalized ATP
125
100
75
50
25
0
500
1,000
5,000
4µm MNP to cell ratio
200µl
500µl
800µl

4μm

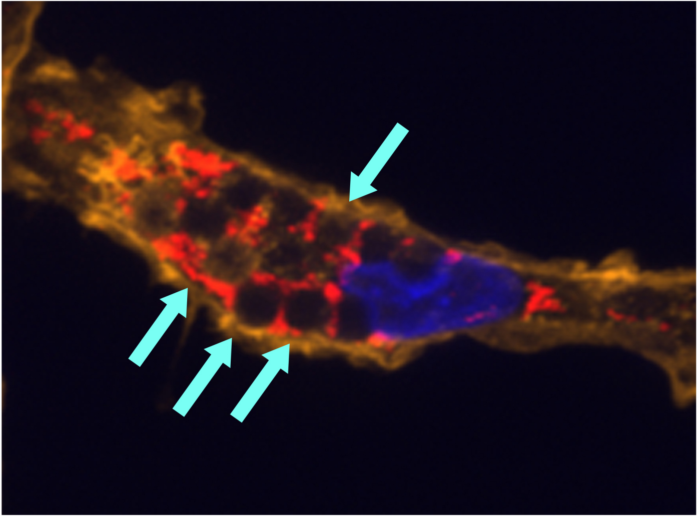

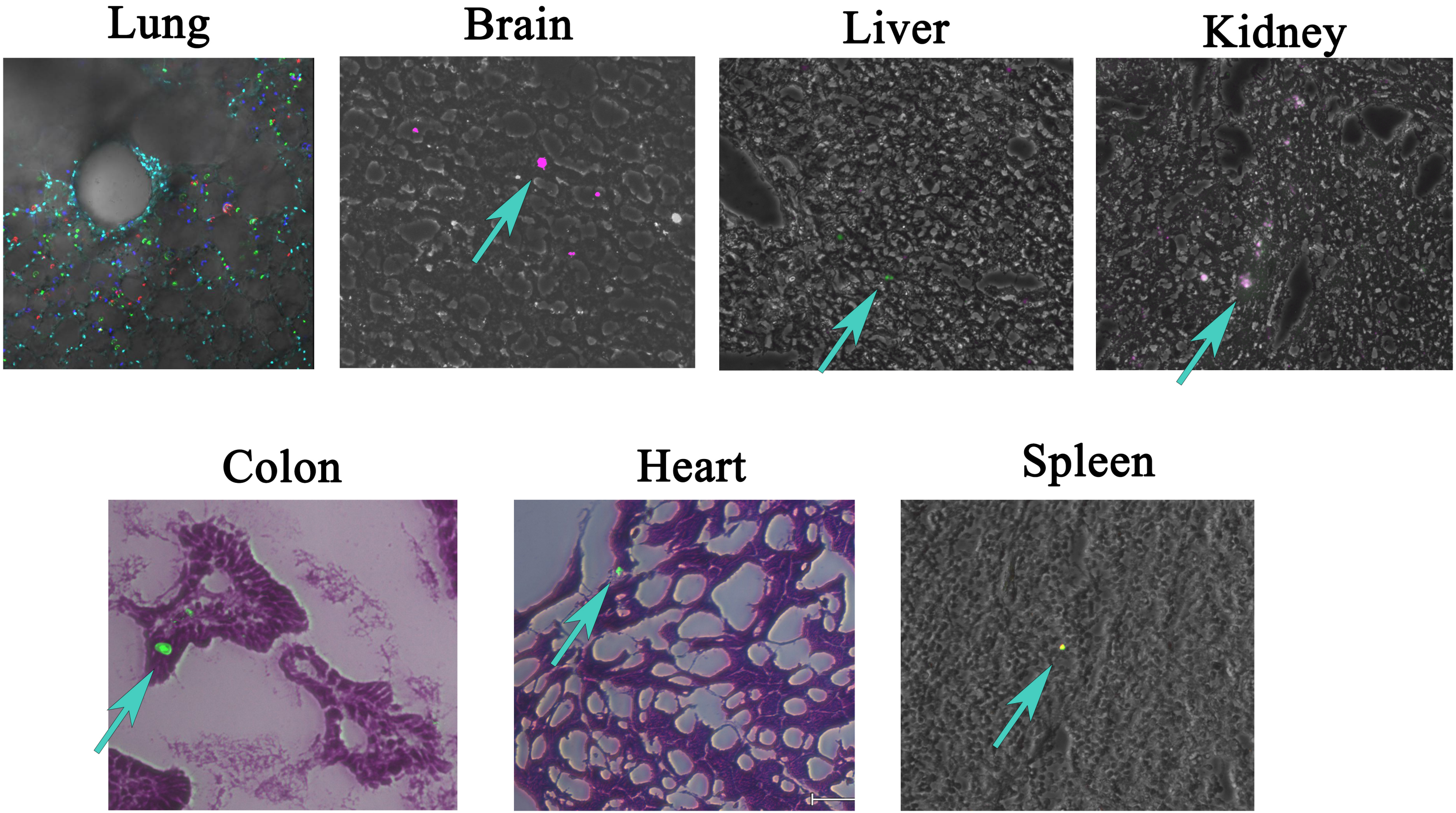

Lung
Brain
Liver
Kidney
Colon
Heart
Spleen
